# A Proteolytic Switch: USP5 controls SDE2 function via UBL-directed cleavage

**DOI:** 10.1101/2025.05.23.655772

**Authors:** Liam T. Hales, Paul M. Tammiste, Adam J. Walker, Rebecca Bryce, Andrew Sugden, Axel Knebel, Rachel Toth, Fred Lamoliatte, Marian Péteri, Oliwia Curry, Virginia De Cesare, Chiara Maniaci

## Abstract

Proteolytic processing is a critical regulatory mechanism in eukaryotic cells, yet the molecular identities and mechanisms underlying these events often remain elusive. Silencing Defective 2 (SDE2) is an essential human protein involved in multiple aspects of genome regulation, including DNA repair, ribosome biogenesis, and mRNA splicing. SDE2 is expressed with an N-terminal ubiquitin-like domain (SDE2_UBL_) which must be proteolytically cleaved to release the functional C-terminal domain (SDE2_CT_). This cleavage not only activates SDE2_CT_ but also marks it for subsequent degradation, highlighting the importance of this tightly regulated processing. Despite the crucial role of this cleavage event, the human protease responsible has remained unknown. Here, we identify that the deubiquitinating enzyme, ubiquitin-specific protease 5 (USP5), catalyses the cleavage of SDE2. Using an integrated workflow combining biochemical assays, proteomic profiling, and mass spectrometry, we demonstrate that USP5 selectively processes SDE2 *in vitro* and in cell. To validate the specificity of this interaction, we engineered SDE2_UBL_ into an activity-based probe, and developed a cellular reporter assay, both of which confirmed USP5 as the primary effector. Biophysical analysis further revealed that SDE2_UBL_ binds to USP5 with similar characteristics to ubiquitin, albeit with reduced affinity, supporting a mechanism of substrate mimicry. Together, these findings uncover a novel regulatory axis for SDE2 function, highlighting an underappreciated role for DUBs in regulating protein maturation events. They also establish a versatile approach for identifying and validating substrate-protease interactions with broader implications for the study of post-translational regulation.

## Introduction

Precise regulation of protein abundance and function *via* insertion or removal of post-translational modifications (PTMs) lies at the heart of cellular homeostasis.^1^ Ubiquitin (Ub) and ubiquitin-like proteins (UBLs) play a key regulatory role by functioning as covalent modifiers that modulate protein activity and direct proteins towards diverse cellular fates, including signalling for degradation, mediating protein-protein interactions, modulating protein activity, and altering protein localisation.^2,3^ These modifications are orchestrated by a sophisticated array of enzymes capable of writing, reading and erasing Ub and UBLs, resulting in a dynamic regulatory system interwoven in nearly every aspect of cell signalling, and therefore often disrupted during pathogenesis.^4,5^ At the centre of this system lies a cascade of enzymatic reactions culminating in the covalent attachment of Ub/UBL to a target protein by an E3 ligase.^6,7^ Antagonising this process are proteases, including deubiquitinating enzymes (DUBs), capable of proteolytically cleaving Ub and UBLs, typically at a conserved C-terminal Gly-Gly motif.^8,9^

While UBLs share high structural similarity with Ub and are often regulated by overlapping enzymatic machinery, many have evolved distinct biological functions through an additional layer of selective interactions with their respective conjugation and deconjugation machinery. For instance, a cascade of interferon induced enzymes, UBEL1-UBE2L6-HECT5, specifically conjugate interferon-stimulated gene 15 (ISG15), rather than Ub, playing an important role in immune response.^10–12^ Similarly, the misleadingly named ubiquitin-specific protein 18 (USP18) specifically cleaves ISG15 from tagged proteins, with no activity toward Ub.^13,14^ These selective interactions facilitate nuanced activation and suppression of Ub and UBL signalling cascades in response to diverse stimuli, and are therefore crucial for effective cellular functioning.

Ub-like domains (ULDs) offer an additional layer of Ub-mimetic regulation without requiring post-translational conjugation.^15^ While Ub acts as a molecular tag by canonically modifying the ε-amino groups of lysine residues, ULDs structurally resemble the Ub fold but are encoded intrinsically within protein sequences.^16^ ULD-bearing proteins often participate in Ub-regulated processes, using the ULD to interact with Ub-binding partners, enabling fine-tuned cross-talk between distinct signalling circuits.^15^ Whilst many ULDs are stable features of a protein’s structure, some contain conserved protease-recognition motifs that permit proteolytic cleavage and irreversible removal of such domains. Proteolysis of such domains resembles the DUB-mediated processing of immature Ub precursors from poly Ub chains (UBB and UBC) or from the ribosomal fusion proteins encoded by the UBA52 and UBA80 genes.^17^ In such cases, these cleavable ULDs act as internally encoded “tags”, with their removal rewiring protein fate and function.

First identified in yeast, Silencing Defective 2 (*H. sapiens*: SDE2, *S. pombe:* Sde2) is one such ULD-containing protein whose proteolytic processing is essential for its biological roles, exemplifying this intriguing mode of regulation.^18,19^ SDE2, a highly conserved and essential gene required for genome maintenance, coordinates essential processes ranging from pre-mRNA splicing and ribosome biogenesis to DNA repair and replication stress response.^18,20,21^ It is expressed as a full-length precursor (SDE2_FL,_ 49.7 kDa) bearing an N-terminal ULD, which is proteolytically cleaved to generate two protein fragments, SDE2_UBL_ (8.35 kDa) and the larger C-terminal domain (SDE2_CT_, 41.4 kDa) (**Figure 1A**). Akin to canonical Ub cleavage, this proteolytic event occurs downstream to a conserved Gly-Gly motif, which becomes exposed following ULD removal, forming the *de novo* C-terminus of SDE2_UBL_.^22^

**Figure 1.**
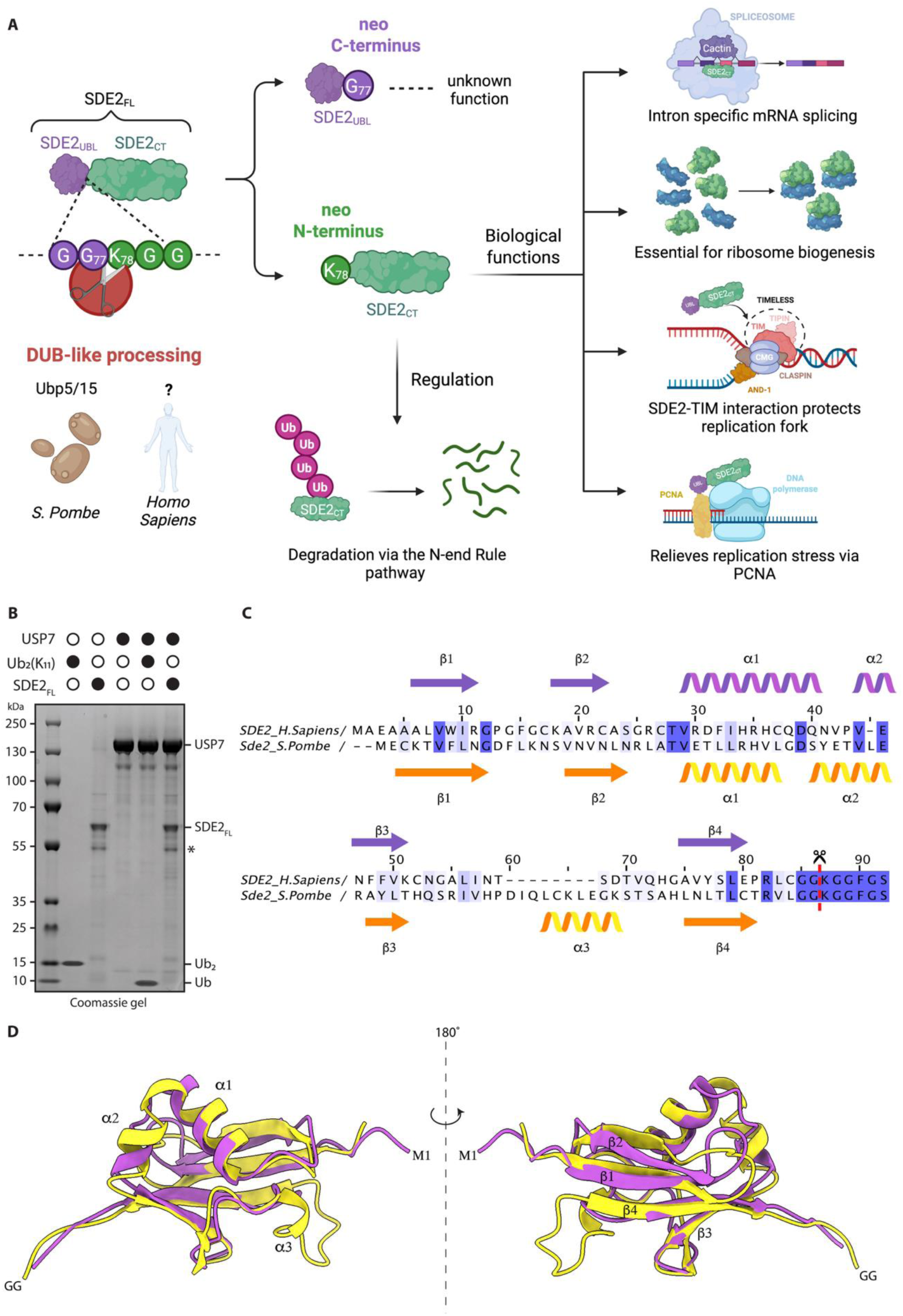
Overview of SDE2 biology and regulation. **(A)** Schematic overview of SDE2 processing, regulation and it’s biological functions. **(B)** USP7 was screened as a potential candidate protease for human SDE2 based on Protein-BLAST analysis of Ubp5/15. USP7 was incubated with either Ub_2_ (K11) or SDE2_FL_ at 37 °C for 1 h and analysed by SDS-PAGE and Coomassie staining. * indicates a protein contaminant. **(C**) Sequence alignment, conservation, and predicted secondary structures of the Ub fold of *H. sapiens* SDE2 (purple) and *S. pombe* Sde2 (yellow/orange). Fully conserved residues are highlighted in dark-blue and semi-conserved residues in light blue. **(D)** Structural comparison of *S. pombe* (yellow) and *H. sapiens* (purple) SDE2 ubiquitin-like fold. Structures were predicted using AlphaFold 3^27^ and visualised in ChimeraX^28^. Secondary structures are highlighted and annotated corresponding to Fig 1C.

The proteolytic cleavage of SDE2 is a crucial regulatory step that underpins its diverse roles in maintaining genome stability, coordinating DNA damage responses, and facilitating proper mRNA splicing. This cleavage is essential for two major functional outcomes. First, the liberated SDE2_CT_, bearing an N-terminal lysine, becomes incorporated into spliceosomes, where it facilitates the efficient excision of specific introns from a subset of pre-mRNAs.^20^ This function is linked to the spliceosomal association of Cactin/Cay1, with genetic and biochemical evidence supporting a functional interplay between SDE2 and Cactin.^20^ Second, the cleaved C-terminal domain interacts with PCNA to negatively regulate damage-inducible monoubiquitination of PCNA, a key modification required for translesion DNA synthesis.^22^ Timely proteasomal degradation of SDE2 is necessary to eliminate this inhibitory activity, thereby permitting S-phase progression and replication fork recovery following DNA damage.^23–25^ Failure to cleave or degrade SDE2 leads to its pathological accumulation, which disrupts the finely tuned balance of PCNA ubiquitination. This dysregulation results in impaired S-phase progression, stalled replication forks, elevated DNA damage, and increased cell death.^20,22^ Thus, SDE2 cleavage serves as a molecular switch that coordinates replication stress responses and RNA processing to safeguard cell proliferation and genome integrity.

Although the regulatory importance of SDE2_UBL_ removal is understood, the enzyme responsible for the cleavage of human SDE2 is unknown. Understanding the mechanisms which control the cleavage of SDE2 and the identification of the protease responsible is crucial to reveal intricacies of SDE2 regulation and the biological functions in which it is involved.

In this study we report the identification of the protease responsible for SDE2 processing, resolving a previously uncharacterised mechanism of post-translational regulation. By developing a targeted approach involving biochemical fractionation of cell lysate coupled with mass spectrometry proteomic analysis, we have identified USP5 as the protease responsible for the cleavage of human SDE2_UBL_, revealing a previously unrecognised, cross-reactive activity of this DUB which extends well beyond its canonical function. Using a combination of tools, including an activity-based SDE2_UBL_ probe and a live-cell BRET reporter assay, we confirm the selective cleavage of SDE2 by USP5 *in vitro* and *in cellulo*. Finally, through biophysical characterisation and mutagenesis studies, we map critical features of USP5 required for this non-canonical recognition event. Together, these findings expand the functional landscape of USP5 and establish a new layer of UBL-driven proteolytic regulation within the cell.

## Results

### Divergence in SDE2_UBL_ processing suggests a distinct human protease

Studies conducted in *S. pombe* identified deubiquitinating enzymes Ubp5 and Ubp15 as the proteases responsible for Sde2_UBL_ cleavage, confirming the role of DUBs in this processing.^20^ This suggested that a related DUB might perform this function in humans. Several human homologues for Ubp15 and 5 have been reported, including ubiquitin-specific proteases (USPs) USP8, USP50, and USP7. We first performed a Protein BLAST analysis^26^ using Ubp5 and Ubp15 as query sequences. The search identified USP7 as the top human match for both yeast enzymes. Despite a modest 34% sequence identity, USP7 shares up to 50% identity with the catalytic domain of Ubp5/15, further supporting its candidacy. To test this hypothesis, we performed an *in vitro* cleavage assay by incubating recombinant USP7 with recombinantly expressed SDE2_FL_. However, while USP7 actively processed Ub_2_ (K11), no processing of recombinant human SDE2 was observed (**Figure 1B**). These findings prompted us to more closely compare full length human SDE2 (451 amino acids) and *S. pombe* Sde2 (263 amino acids) to understand potential structural divergence. Despite a 39% sequence identity, conservation is concentrated around the GGKGG cleavage motif (**Figure 1C and S1**), suggesting a preserved recognition element. Secondary structure prediction of SDE2_UBL_ and Sde2_UBL_ revealed strong alignment between the two UBL domains, with the major difference being a loop and alpha helix (α3) (amino acids D59-E65) present in the yeast protein but absent in the human SDE2 (**Figure 1C**). Structural modelling using AlphaFold 3^27^ confirmed the high structural similarity of the UBL folds (RMSD = 1.574). However, the model also highlighted a loop and alpha helix (α3) as divergent structural elements positioned near the N-terminal tail, distant from the conserved C-terminal cleavage site (**Figure 1D**). These observations indicate that, despite structural conservation in the cleavage region, species specific differences may influence protease recognition. The failure of USP7 to cleave SDE2_UBL_ *in vitro* suggests that the human protease responsible may differ substantially from those in yeast, either in sequence or in mechanism of substrate engagement. As such, identifying the human protease responsible for SDE2_UBL_ cleavage is critical to understanding how SDE2 is regulated, and may reveal novel principles of substrate recognition by DUBs.

### Integrated biochemical and proteomic strategy implicates USP5 and USP13 in SDE2 proteolysis

To uncover the identity of the protease responsible for processing SDE2 in human cells, we first turned to a biochemical strategy using recombinant SDE2_FL_ as substrate (**Figure 2A**). Recognising that protease activity can be sensitive to buffer conditions, we lysed Expi293 cells in hypotonic conditions specifically chosen to preserve endogenous enzymatic function. The crude lysate was then subjected to a two-step column chromatography workflow (**Figure 2A**). In the first step, we applied the lysate to a heparin column, to remove nucleic acid-binding and regulatory proteins, followed by ion-exchange chromatography using a Source-Q column to separate proteins based on charge. This approach resolved the lysate into discrete fractions while maintaining enzymatic integrity. Next, each collected fraction was incubated with recombinant SDE2_FL,_ and proteolytic activity assessed by monitoring the formation of the expected cleavage products, SDE2_CT_ and SDE2_UBL_ fragments, observed *via* SDS-PAGE (**Figure 2A**). Proteolytic activity was highly restricted to two adjacent SourceQ column fractions, A12 and B1, indicating a successful enrichment of candidate protease (**Figure 2B**). To further isolate the active enzyme, active fractions were further purified through size exclusion chromatography (SEC), reasoning that this additional resolution step would help distinguish true proteolytic activity from co-eluting contaminants. SEC fractions were once again tested for their ability to cleave recombinant SDE2_FL_, revealing that activity was concentrated in fractions B2 through B4 (**Figure 2C**). These fractions, as well as surrounding ones (A11-B1 and B5-B7), were subjected to tryptic digestion followed by mass spectrometry. Proteomic analysis identified 24 proteins common to the active fractions. Among these, only two proteins, USP5 and USP13, were annotated with known or predicted protease activity (**Figure 2D**). This finding was particularly intriguing as it aligned with the precedent reports of DUB-mediated Sde2 processing in *S.pombe*.^20^ Western blot analysis confirmed the presence of both USP5 and USP13 across the active fractions from both Source-Q and SEC chromatography steps, strengthening the case for their involvement (**Figure 2E**). To rigorously test whether these DUBs could indeed cleave SDE2, we conducted an orthogonal screening assay using a panel of 70 recombinant human and viral proteases, including 65 DUBs. Each enzyme was incubated individually with recombinant SDE2_FL_, and cleavage was assessed by MALDI-TOF/MS, focusing on detection of SDE2_UBL_ as the readout of activity (**Figure 2F**). Among the entire panel, only USP5 and USP13 consistently cleavedSDE2_FL_ across biological replicates. Notably, USP5 exhibited the highest level of activity cleaving approximately 50% of the input substrate under the assay conditions (**Figure 2G**). Together, these results implicate USP5 and USP13 as leading candidates for the processing of human SDE2, with USP5 emerging as the dominant enzyme in recombinant system.

**Figure 2.**
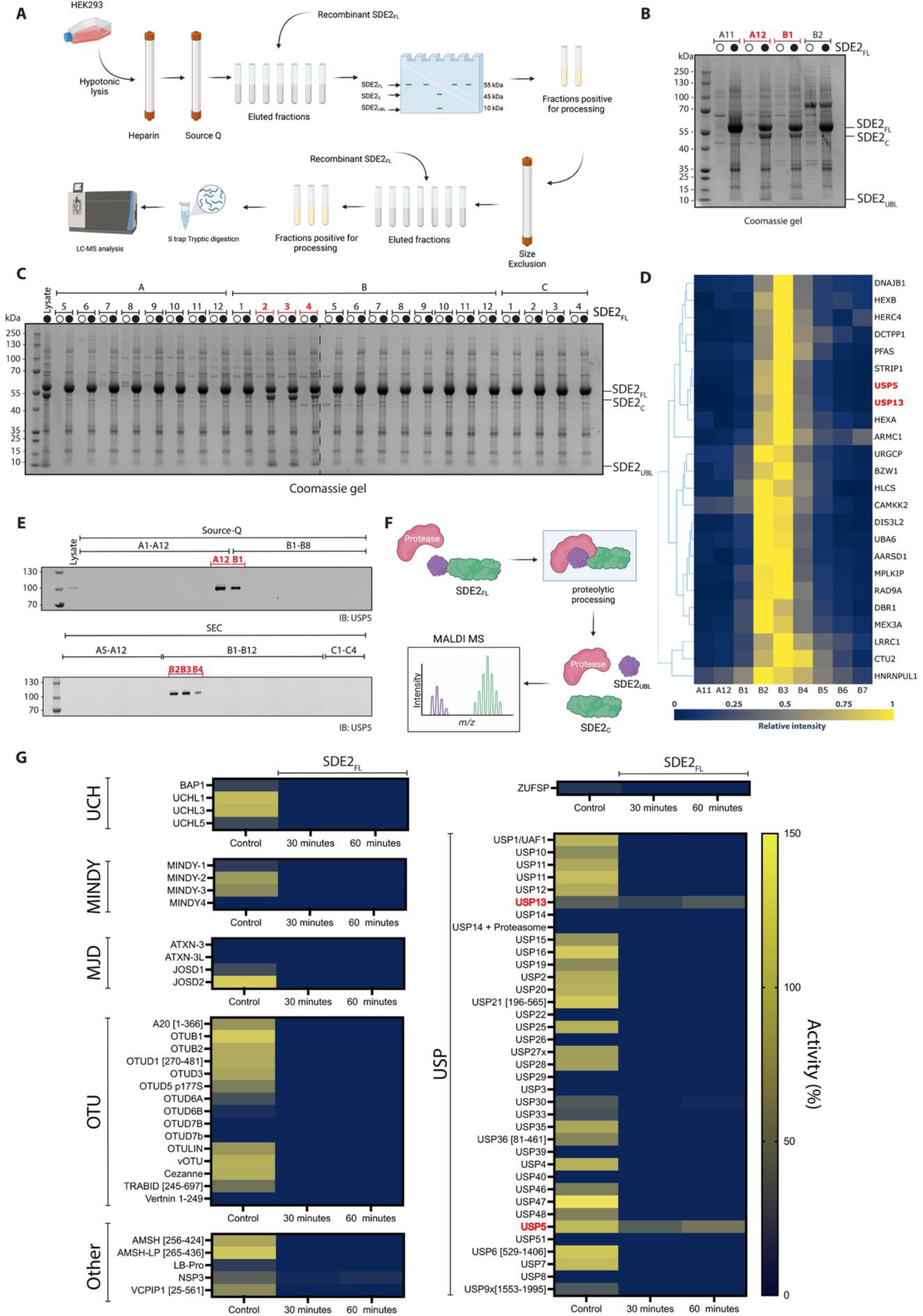
Screening for SDE2 protease identified USP5 and USP13 as candidate proteases. **(A)** Schematic overview of the lysate screening process. Expi293 cells were lysed in hypotonic conditions and fractionated sequentially over Heparin and Source-Q columns. Eluted fractions were screened for activity by incubation with recombinant SDE2_FL_. Fractions showing activity were by through Size Exclusion Chromatography (SEC) and collected fractions were screened again for processing. Active fractions were subjected to tryptic digestion over S-trap column and analysed by mass spectrometry. **(B)** Representative screening results for fractions collected after Source-Q ion-exchange chromatography. Eluted fractions were incubated with SDE2_FL_ (black dot) for 30 min at 37 °C and analysed by SDS-PAGE and Coomassie staining. Protein cleavage activity was detected in fractions A12 and B1 (red lines). **(C)** Representative screening results for A12 and B1 fractions collected after SEC. Proteins were eluted in fractions A1-C4. Eluted fractions were incubated with SDE2_FL_ (black dot) for 30 min at 30 °C and analysed by SDS-PAGE and Coomassie staining. Protein cleavage activity was detected in fractions B2-B4 (orange lines). **(D)** Identity of proteins in all active fractions as detected by proteomic mass spectrometry. **(E)** Immunoblot analysis of endogenous USP5 in fractions from Source-Q and SEC. Fractions that showed activity are highlighted in orange. **(F)** Schematic overview of the high-throughput MALDI-TOF SDE2 cleavage assay. Recombinant proteases were incubated with SDE2_FL_ and the amount of SDE2_UBL_ formed by this reaction was determined by MALDI-TOF MS by the presence of peak for SDE2_UBL_ at *m/z*= 8348.9. **(G)** 70 enzymes including known DUBs, pseudo-DUBs and proteases were incubated with their respective specific substrates (control) or with SDE2_FL_ for 30 or 60 min at 30 °C and analysed by the MALDI-TOF MS. The amount of SDE2_UBL_ formed by this reaction was determined by MALDI-TOF/MS and quantified against ^15^N-Ub internal standard. Activity for the individual tested enzymes is reported in a gradient of blue (0%) to yellow (150%).

### An SDE2_UBL_-derived activity-based probe reacts with USP5, but not USP13, in a catalytically dependent manner

Having identified USP5 and USP13 as the candidate proteases responsible for cleavage of SDE2_UBL_, we next focused on identifying the functional amino acids involved in the SDE2_FL_ cleavage event. To do this, we focused on the region of 20 amino acid between S69 and L88, including the previously reported cleavage site motif GGK.^20,22^ We employed an alanine scanning approach where we generated a library of SDE2 expressing plasmids where every amino acid in the selected region was individually mutated to alanine. As controls, we also included a plasmid expressing SDE2_FL_ wild type (WT) and a G76A G77A double mutant, previously reported to abrogate processing.^20,22^ HEK293T cells were transiently transfected with each individual plasmid and SDE2 processing was evaluated by western blot. Interestingly, the mutations that impaired SDE2 processing were all localised in the region of UBL domain. Indeed, while none of the tested mutations in the region corresponding to the start of SDE2_CT_ seem to affect SDE2 cleavage, several mutations in the SDE2_UBL_ region seem to reduce or abrogate SDE2 processing to different degrees. As expected, mutagenesis of the GG motif, both single and double mutations, resulted in impairment of SDE2 processing (**Figure 3A**). Mutations L70A and R73A also completely prevented SDE2 cleavage. Mutations of amino acids E71 and L74 resulted in around a 50% cleavage of SDE2 suggesting that these amino acids play a supporting role in the cleavage process. These results suggest that the SDE2_UBL_-USP5/USP13 interaction could be crucial driving force for the proteolysis.

**Figure 3.**
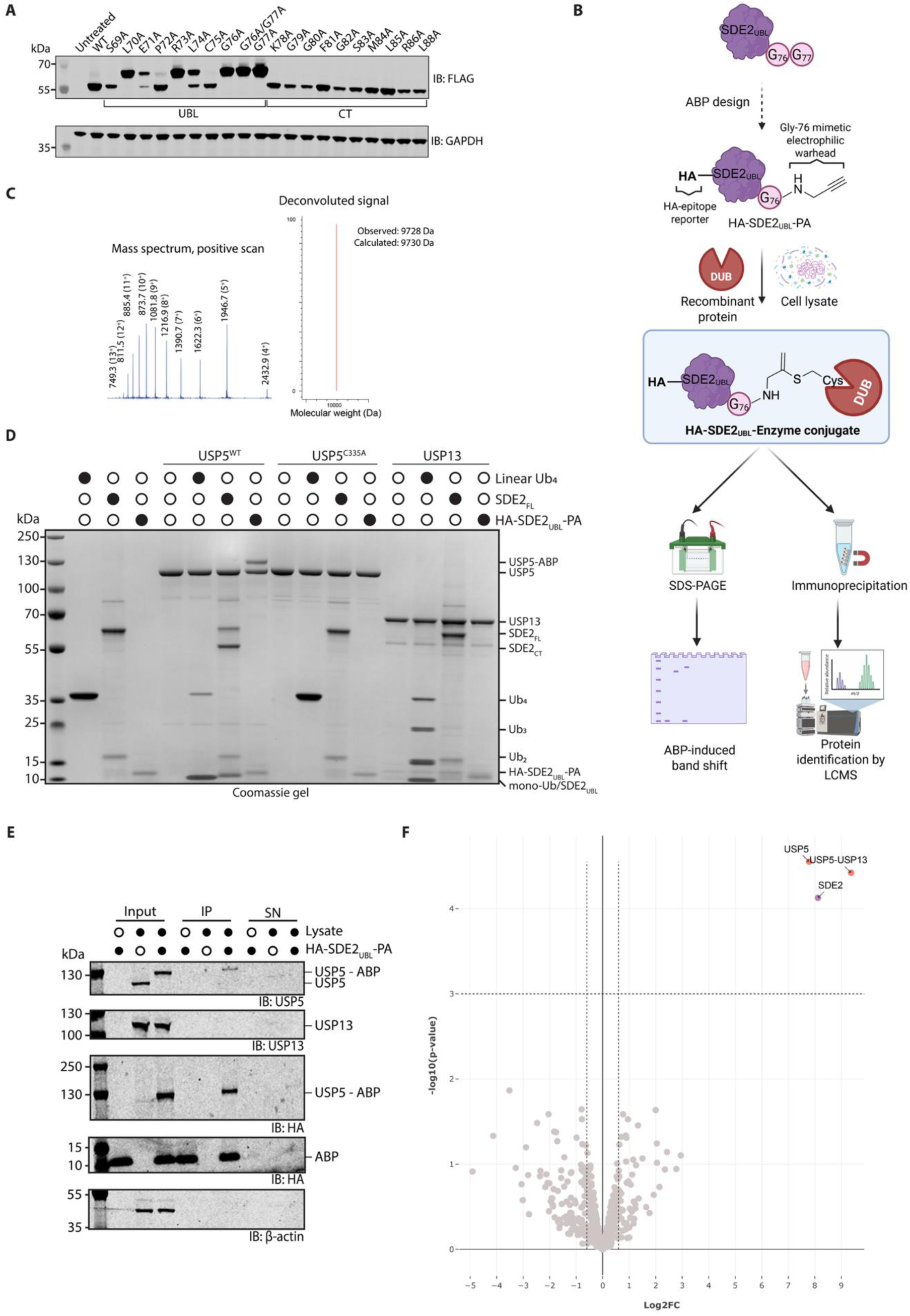
An ABP derived from SDE2_UBL_ reacts selectively with USP5, but not USP13, in *in vitro* reconstitution assay and cell lysate contexts. **(A)** Representative western blot results from alanine scanning of the residues adjacent to the site of cleavage. HEK293T cells overexpressing HA-SDE2-FLAG WT or indicated mutants were lysed and immunoblotted for the FLAG epitope to determine the effect of each mutation on SDE2_FL_ cleavage. The location of the residues is annotated as either the UBL or C-terminal domain (CTD). **(B)** An ABP derived from SDE2_UBL_ was designed and prepared semi-synthetically. Gly77 is replaced with a propargyl electrophilic warhead to mimic the scissile amide bond. An N-terminal HA epitope tag facilitates recognition by immunostaining. This ABP can be used to detect catalytic-dependent interactions with enzymes, either by incubation with recombinant protein (visualisation by SDS-PAGE), or with cell lysate followed by HA-mediated immunoprecipitation and analysis by western blot and mass spectrometry. **(C)** Mass spectrum of HA-SDE2_UBL_-PA obtained using intact LCMS. **(D)** Assessment of USP5^WT^/USP5^C335A^ and USP13 reactivity towards various substrates. DUBs (1 µM) were incubated with HA-SDE2_UBL_-PA (2 µM), SDE2_FL_ (2 µM) or linear tetraubiquitin (Ub_4_, 1 µM) for 1 h at 37 °C and analysed by SDS-PAGE. (**E)** Immunoblot analysis following immunoprecipitation of HEK293T lysate treated with HA-SDE2_UBL_-PA. HEK293T cell lysate was incubated with HA-SDE2_UBL_-PA (1 µM) for 45 min at 37 °C and immunoprecipitated with anti-HA beads. **(G)** Volcano plot analysis of enriched HEK293T cell lysate following HA-SDE2_UBL_-PA treatment. ABP immunoprecipitated samples (triplicate) were subject to trypsin digestion and proteomic analysis, including comparison to the UniProt Swiss-Prot human database to identify the parent protein. Samples were compared to a ‘no-probe’ negative control.

Given the importance of the UBL domain sequence for SDE2_FL,_ we next sought to validate the SDE2-USP5/USP13 interaction through the development of an SDE2_UBL_ activity-based probe (ABP). We designed a semi-synthetic approach for the synthesis of a hemagglutinin (HA)-tagged SDE2_UBL_ probe. Here, SDE2_UBL_ serves as a recognition subunit and the N-terminal HA tag serves as both an enrichment handle and a reporter tag for easy detection via immunoblotting, allowing us to profile protease enzymatic activity in both recombinant and complex biological mixtures (**Figure 3B**). Many UBLs have been converted into ABPs through the installation of electrophilic warheads which mimic the C-terminal scissile amide bond. For the majority of UBLs, this involves the replacement of the terminal glycine with an electrophilic glycine mimetic. This approach has been successfully employed for the activity-based profiling, aiding in the identification of novel DUBs,^29^ or of UBLs as novel substrates for various DUBs.^30^ In this case, a propargylamine warhead was installed as covalent trap for active SDE2-interacting cysteine enzymes.

To generate a functional SDE2_UBL_ probe, we employed a semisynthetic approach by leveraging the intein-mediated ligation method, a method widely used to generate UBL ABPs semi-synthetically.^31^ An N-terminally HA-tagged SDE2_UBL_ containing a Gly77 truncation (HA-SDE2_UBL_ΔG77) was expressed as a fusion with a tandem self-cleaving intein and chitin-binding domain, allowing for efficient purification using chitin resin. Cleavage from the resin was achieved by addition of 2-mercaptoethane sulfonate (MESNA), which yielded a reactive thioester at its C-terminus (HA-SDE2_UBL_-MESNA). The final ABP (HA-SDE2_UBL_-PA, **Figure 3B**) was generated by adding excess propargylamine to initiate the nucleophilic substitution of the thioester.

A propargyl warhead was selected due to its unique electrophilic characteristics which prevent it from reacting with untargeted cysteines (Cys), minimizing non-specific interactions with the Cys present in SDE2_UBL_ sequence.^32,33^ Indeed, unlike other UBLs which either lack Cys or contain a single Cys residue which can be easily mutated, SDE2_UBL_ contains six Cys residues in its sequence. As a result, attempts to utilise more reactive electrophiles were unsuccessful. Notably, mass spectrometry revealed evidence of MESNA conjugation to these native cysteines following cleavage from the chitin resin, indicated by mass shifts consistent with one or more MESNA adducts (**Figure S2**). These adducts were successfully removed by DTT treatment following propargylamine conjugation, restoring the expected ABP mass (**Figure 3C**).

To assess the probe reactivity, we incubated HA-SDE2_UBL_-ABP with recombinant wild-type USP5 (USP5^WT^), catalytically inactive USP5 and USP13. HA-SDE2_UBL_-ABP readily reacted with USP5^WT^, evidenced by the formation of a higher molecular weight band by SDS-PAGE analysis, corresponding to the covalently linked ABP-USP5 adduct (**Figure 3C**). The catalytically inactive USP5^C335A^ mutant neither reacted with the ABP nor cleaved SDE2_FL_, indicating that both activities are dependent on the cysteine mediated protease activity of USP5. In contrast, USP13 showed no evidence of probe labelling, suggesting that the cleavage of SDE2_FL_ observed in the active SEC fractions is primarily driven by USP5.

Next, to exclude the possibility that USP13 may require a cofactor for activity, we utilised the ABP to perform activity-based profiling in HEK293 cell lysate. Incubation of the lysate with the SDE2_UBL_-ABP followed by HA immunoprecipitation allowed us to isolate covalently labelled DUBs (**Figure 3E**). Immunoblotting with a specific α-USP5 antibody revealed a distinct band shift upon ABP treatment, indicating near complete labelling of endogenous USP5. The shifted band was also detected with α-HA, confirming it represented the HA-SDE2_UBL_-USP5 complex. No labelling of USP13 was observed, consistent with our recombinant protein data and previously reported literature.^34^

To further validate the identity of the probe-bound proteins, we performed a tryptic digestion and mass spectrometry analysis on the ABP-enriched material. Proteomics analysis revealed strong enrichment of peptides specific for USP5 and SDE2, confirming the formation of a covalent HA-SDE2_UBL_-USP5complex (**Figure 3F**). Although some USP13-related peptides were detected, these corresponded to regions of shared homology with USP5. Critically, no USP13-specific peptides were identified, reinforcing the conclusion that USP5—but not USP13—directly interacts with SDE2_UBL_ in cells. Furthermore, proteomic analysis did not reveal binding of the probe to other DUBs, indicating a high level of selectivity for USP5. Together, these results provide strong biochemical and cellular evidence that USP5 is the primary protease responsible for SDE2_UBL_ cleavage, and they establish HA-SDE2_UBL_-PA as a selective and effective probe for USP5 activity.

### Knockdown cell assay reveals that SDE2 is selectively processed by USP5

To determine whether the proteolytic selectivity observed *in vitro* extends to a cellular context, we developed a sensitive, real-time assay to monitor SDE2 processing in live cells using the bioluminescence resonance energy transfer (NanoBRET) technology.^35^ In this system, SDE2 was overexpressed as a fusion protein with an N-terminal NanoLuc luciferase tag, acting as an energy donor, and a C-terminal Halo-Tag as an energy acceptor. When intact, for example in the case of impaired processing, upon treatment with a fluorophore-containing chloroalkene ligand, the proximity of the donor and acceptor allows for efficient dipole-dipole energy transfer from the luciferase (NanoLuc) to a fluorescently labelled HaloTag, resulting in a detectable BRET signal. Upon proteolytic cleavage of SDE2_FL_, the donor and acceptor are separated, leading to a measurable loss of energy transfer and thus a decrease in BRET signal (**Figure 4A**).

**Figure 4.**
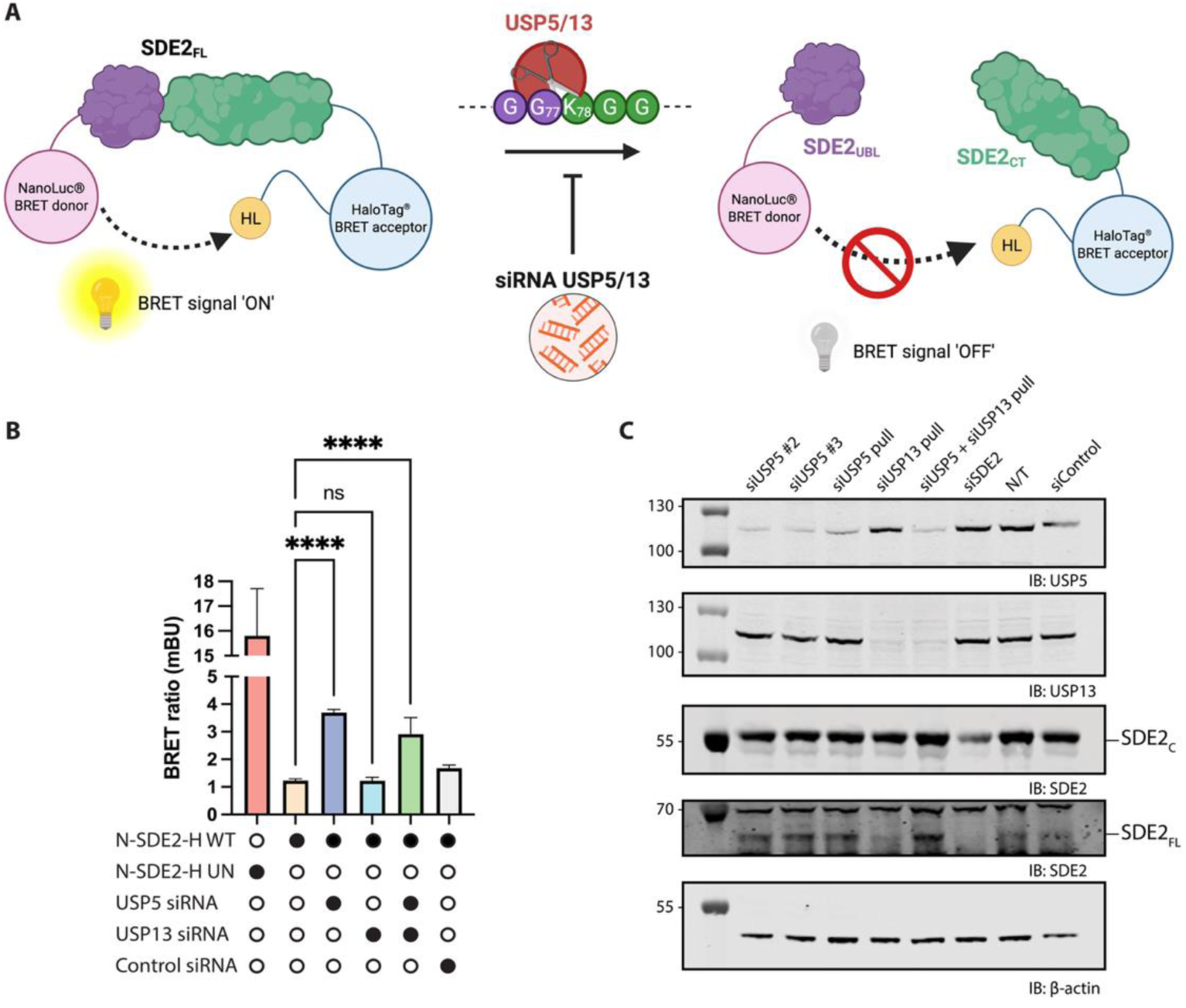
siRNA knockdown confirms USP5 as solely responsible for SDE2 processing at both overexpression and endogenous levels. **(A)** Schematic representation of a NanoBRET intracellular SDE2 cleavage assay. SDE2 is tagged with an N-terminal NanoLuc (N-SDE2, donor) and a C-terminal HaloTag (SDE2-H, acceptor). Upon addition of HaloTag 618 Ligand, if SDE2 is processed, the SDE2_UBL_ and SDE2_CT_ fragments will separate, and no BRET signal will be detected. If the processing is impaired, the donor and acceptor tags will be in close proximity, so the dipole-dipole non-radiative energy transfer from the luciferase energy donor (NanoLuc) to the acceptor fluorophore (HaloTag-bound fluorescently labelled chloroalkene ligand) will lead to emission of a light signal. **(B)** Calculated NanoBret ratio in experimental samples. WT= wild-type, UN= uncleavable. Graphs depict the mean + SEM of three independent biological replicates. One way ANOVA was performed to calculate P values, and levels of significance are denoted as follows: ****P<0.0001, ns=not significant. **(C)** U2OS cells were treated with siRNA targeting USP5, USP13, SDE2 or control siRNA for 48h. A not treated control (NT) was also included. The data is a representative example of experiments performed as three biological replicates.

To validate the system and provide a positive control, we engineered an uncleavable (UN) SDE2 variant in which the conserved cleavage motif -GGK-motif was mutated to -AAK-ensuring that the donor and acceptor remain in close proximity and the BRET signal remains constant and high, allowing highly specific, quantitative monitoring of SDE2 cleavage dynamics in live cells.

Using this assay, we tested the contribution of USP5 and USP13 to SDE2 processing *via* siRNA-mediated knock down (**Figure 4B and S3**). Depletion of USP5, either alone or in combination with USP13, resulted in a significant increase in BRET signal (****P<0.0001). In contrast, USP13 knockdown alone had no significant effect, suggesting it does not play a major role in SDE2 processing under these conditions. These findings support the conclusion that USP5 is the primary protease responsible for SDE2_UBL_ cleavage in cells, and that it can function independently of USP13.

To ensure that the observed effects were not an artifact of SDE2 overexpression, we also performed immunoblot analysis of endogenous SDE2 levels in cells treated with siRNA targeting USP5, USP13 or both (**Figure 4C**). In these experiments, knockdown of USP5 using three distinct siRNA constructs consistently impaired SDE2 processing, as indicated by the accumulation of SDE2_FL_. In contrast, USP13 knockdown did not affect SDE2 cleavage, and only cells treated with USP5-targeting siRNA or combined USP5/USP13 knockdown exhibited robust rescue of SDE2_FL_, further confirming that USP5 is solely responsible for SDE2_UBL_ cleavage in cell.

### Reactivity of USP5 towards SDE2_UBL,_ Ub and ISG15 ABPs reflects preferential processing of substrates

Canonically, USP5 is known for its role in processing unanchored polyubiquitin chains, a function that is critical for maintaining the pool of free Ub and ensuring proper Ub signalling dynamics. Beyond this activity, USP5 can also deconjugate Ub from modified substrates and, as expected, shows strong reactivity with Ub-based ABPs.^36,37^ Notably, prior studies have also reported USP5 reactivity for ISG15-derived ABPs.^30,38,39^ This cross reactivity suggests that USP5 may exhibit broader UBL recognition than previously appreciated.

To systematically assess USP5 reactivity, we performed a comparative labelling assay using three ABPs, Ub-PA, ISG15-PA and HA-SDE2_UBL_-PA. We incubated these ABPs with USP5, alongside the prototypical DUB, USP7, and a panel of DUBs previously reported to react with ISG15-PA, enabling a direct comparison of DUB reactivity (**Figure 5A**).^40,41^ All tested DUBs reacted with Ub-PA, confirming their enzymatic competence. HA-SDE2_UBL_-PA displayed a high degree of selectivity, exclusively labelling USP5, in agreement with our previous immunoprecipitation-MS data (**Figure 3F**). In contrast, ISG15-PA exhibited broader promiscuity, covalently modifying USP5, USP16 and USP21. Together, these data highlight the unique reactivity profile exhibited by USP5 towards these UBL-ABPs.

**Figure 5.**
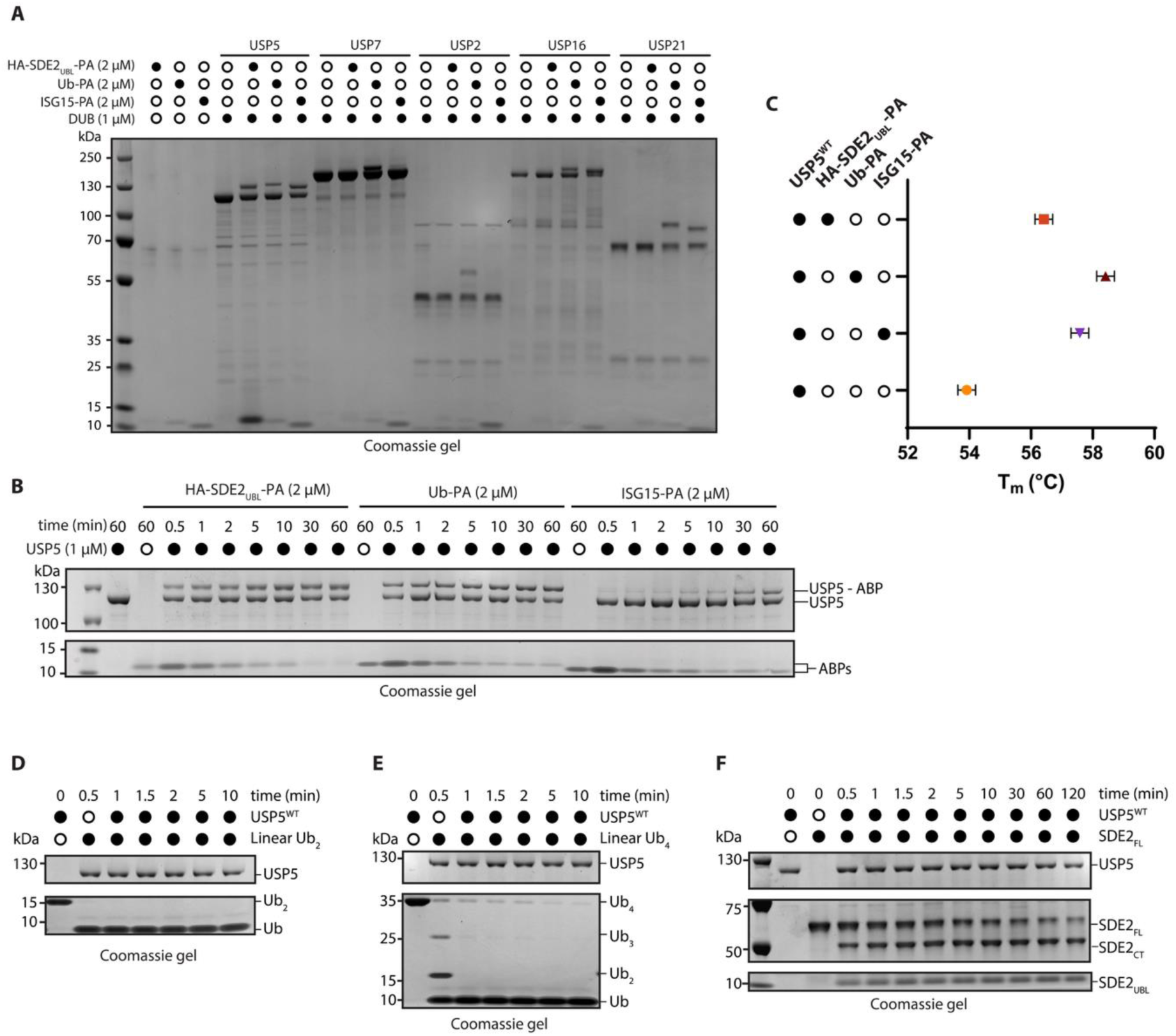
USP5 exhibits a preference for Ub based substrates. **(A)** The reactivities of HA-SDE2_UBL_-PA, Ub-PA or ISG15-PA were compared across a panel of DUBs. Each DUB (1 µM) was incubated separately with each ABP (2 µM each) for 1 h at 37 °C. A band shift was observed for ABP-reactive DUBs by SDS-PAGE and Coomassie staining. **(B)** Time course assay to assess the kinetic of ABP reactivity. USP5^WT^ (1 µM) was incubated separately with each ABP for 1 h at 37 °C with a sample taken at the indicated time points. **(C)** Thermal shift analysis of USP5-ABP complexes. USP5^WT^ (1 µM) was incubated with each ABP for 1 h at 37 °C before thermal cycling from 10-95 °C in the presence of SYPRO orange. Calculated melting temperatures (T_m_) indicate the stability of the USP5-ABP covalent complex. **(D-F)** Comparison of USP5-mediated cleavage of SDE2_FL_ and Ub substrates. USP5^WT^ (1 µM) was incubated at 37 °C with linear diubiquitin (Ub_2_, D, 3 µM), linear tetraubiquitin (Ub_4_, E, 3 µM), or SDE2_FL_ (F, 3 µM). Samples were taken at the indicated time points and cleavage assessed by SDS-PAGE and Coomassie staining.

We next compared the kinetics of covalent adduct formation between these ABP and USP5. Both Ub-PA and HA-SDE2_UBL_-PA exhibited rapid labelling kinetics (**Figure 5B**), with detectable USP5 modification evident after 30 sec and near-complete labelling observed between 10 and 30 min. ISG15-PA, by contrast, labelled USP5 much more slowly, with a significant portion of unmodified USP5 persistent even after 60 min incubation. Despite its slower labelling kinetics, ISG15-PA formed a more thermally stable complex with USP5, as evidenced by a thermal shift assay. The melting temperature (T_m_) of the USP5-ISG15-PA complex was higher T_m_ than that of USP5-SDE2_UBL_-PA complex (57.6 °C ± 0.2 vs 56.4 °C ± 0.2, respectively, **Figure 5C**).

Considering the preferential reactivity towards HA-SDE2_UBL_-PA and Ub-PA, we next compared the processing of the substrates from which these ABPs are derived. A gel-based cleavage assay using recombinant SDE2_FL_, alongside linear di- and tetraubiquitin substrates revealed that USP5 demonstrated a clear preference for Ub substrates, achieving near complete processing of both substrates after 120 sec (**Figure 5D/E**). In contrast, cleavage of SDE2_FL_ was slower and incomplete even after 2 hours, suggesting distinct substrate engagement dynamics (**Figure 5F)**. Notably, recombinant SDE2_FL_ processing appeared to stall after 30 min, in contrast to the complete processing observed for cellular SDE2 (**Figure 3A**).

Collectively, these data underscore both the versatility and unique reactivity of USP5. These findings highlight the nuanced molecular interactions that govern USP5 activity and suggest that substrate context plays a pivotal role in modulating USP5 engagement and catalysis.

### Selectivity and affinity for SDE2_UBL_ drives the selective interaction between SDE2_FL_ and USP5

While HA-SDE2_UBL_-PA is a valuable tool for activity-based profiling, it relies on covalent modification and may not fully reflect native binding interactions. To better understand the native interaction between SDE2 to USP5, we used isothermal titration calorimetry (ITC). Using catalytically inactive USP5^C335A^ to prevent cleavage, we compared the interaction of SDE2_FL_ with that of the individual domains, SDE2_UBL_ and SDE2_CT_. Among the constructs tested, SDE2_UBL_ bound to USP5^C335A^ with the highest affinity (*K*_d_ = 1.67 ± 0.62 µM, **Figure 6A**), consistent with a specific recognition interface. By contrast, SDE2_FL_ showed only weak, non-quantifiable binding (**Figure 6B**). Titration of the SDE2_CT_ resulted in no detectable interaction (**Figure 6C**), indicating that this domain neither contributes to, nor enhances binding. These findings suggest that SDE2_CT_ does not contribute crucial additional interactions with USP5.

**Figure 6.**
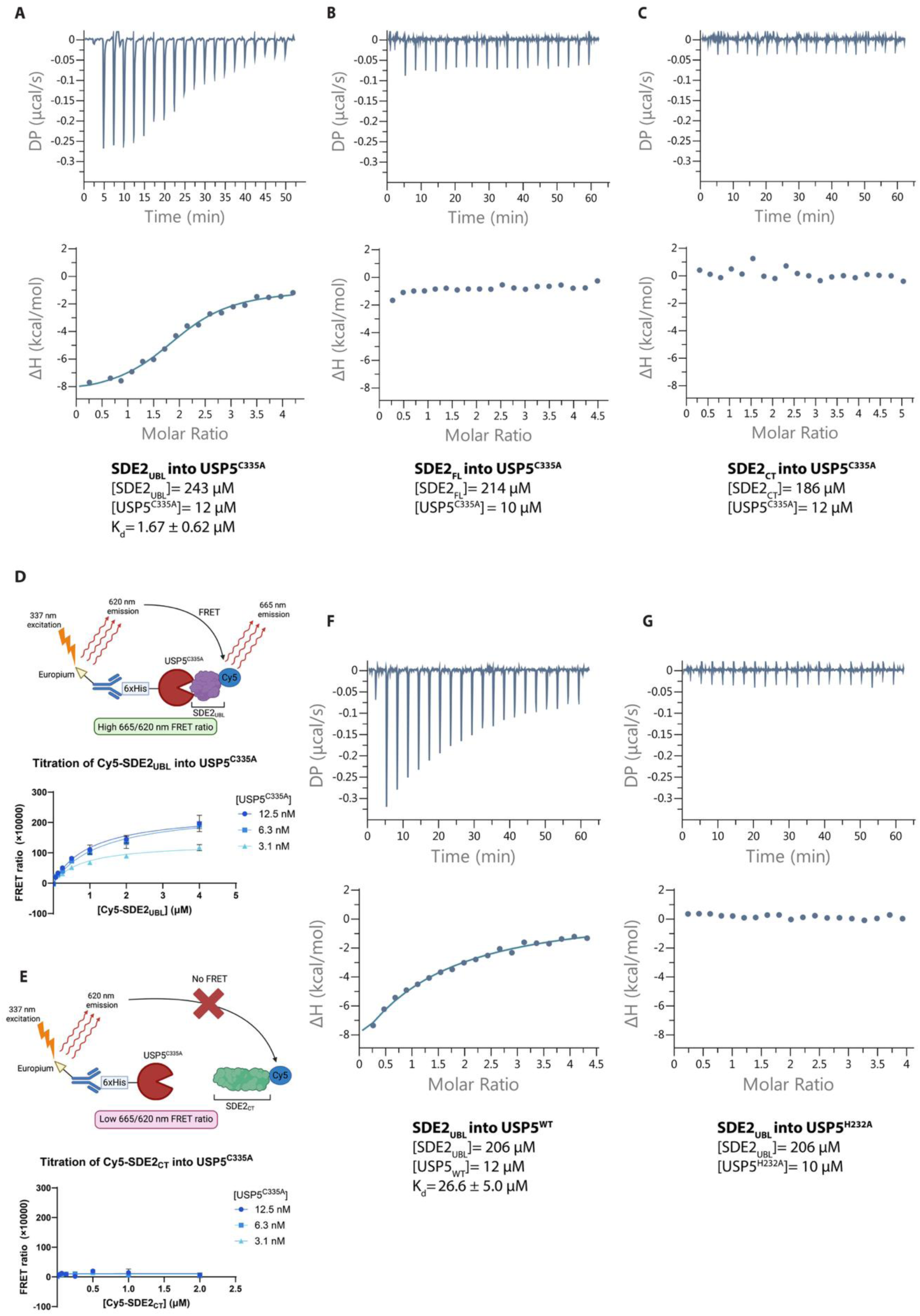
Assessment of binding between USP5^C335A^ and SDE2 constructs confirms that SDE2_UBL_ is the primary driver of the interaction. **(A-C)** ITC analysis of binding between USP5 and SDE2 constructs. SDE2_UBL_ (A, 243 µM), SDE2_FL_ (B, 206 µM) or SDE2_CT_ (C, 186 µM) were titrated into USP5^C335A^ (10-12 µM). Isotherm curves are representative of experiments performed in duplicate. The *K*_d_ is presented as the mean + SEM of the two experiments. **(D-E)** A TR-FRET assay was established to monitor binding between USP5 and SDE2 domains. His_6_-tagged USP5 is preincubated with an Eu-tagged His_6_-antibody (FRET donor), followed by the addition of a Cy5-labelled SDE2 construct (FRET acceptor). The ratio of light emitted at 665 nm to 620 nm is calculated to quantify binding affinity. Cy5-SDE2_UBL_ (D) or Cy5-SDE2_UBL_ (E) was titrated into USP5^C335A^ (at 3.1, 6.3 or 12.5 nM) and the FRET ratio was measured. An increasing FRET ratio indicative of binding was observed upon addition of Cy5-SDE2_UBL_, but not Cy5-SDE2_CT_. TR-FRET experiments were performed as technical triplicates and curves fitted using a one-site specific curve to calculate a *K*_d_. **(H-I)** ITC analysis of SDE2_UBL_ binding to USP5^WT^ and USP5^H232A^ confirms that SDE2_UBL_ occupies a similar binding site to Ub. SDE2_UBL_ (206 µM) was titrated into USP5^WT^ (F, 12 µM) or USP5^H232A^ (G, 10 µM).

To further validate this interaction, we developed a time resolved fluorescence energy transfer (TR-FRET) assay. Cy5 labelled SDE2_UBL_ or SDE2_CT_ were titrated into His-USP5^C335A^ in presence of europium (Eu) labelled α-His antibody, leveraging a proximity dependent FRET interaction between Cy5 and Eu to detect binding (**Figure 6D**). Consistent with the ITC data, Cy5-SDE2_UBL_ showed robust binding to USP5^C335A^ with *K*_d_= 1.0 ± 0.1 µM (**Figure 6E**), while Cy5-SDE2_CT_ exhibited no appreciable interaction (**Figure 6F**). Titration of Cy5-SDE2_FL_ resulted in flat binding curves (**Figure S4**). Together, these data confirm that USP5 specifically engages the SDE2_UBL_ domain and has no intrinsic affinity for SDE2_CT_.

Given this specificity, we next sought to define the molecular determinates of SDE2_UBL_ recognition by USP5, particularly in relation to known Ub-binding features as well as understanding the importance of the characteristic zinc finger ubiquitin-binding domain (ZnF-UBP) domain present in the structure of USP5. USP5 contains four Ub binding sites, facilitating tetraubiquitin processing. These sites are composed of the ZnF-UBP, USP, UBA2 and UBA1 domains (**Figure S5A**), and within each site residues have been identified which are essential for Ub binding.^42–44^ The ZnF-UBP can be disrupted by a H232A mutation, removing the His residue essential for Zn^2+^ coordination. Mutation of methionine 743 (UBA2, S2 site) or 666 (UBA1, S3 site) to glutamic acid disrupts key hydrophobic interactions. Binding of Ub to the USP site, housing the catalytic triad, is disrupted by a C335R mutation which serves as an alternative to the standard C335A catalytically inactive mutant.

Comparison of Ub and SDE2_UBL_ binding to USP5, as predicted by AlphaFold3^27^, suggests similar binding modes, with both UBLs placed within the catalytic site of USP5 (**Figure S5B**). We therefore hypothesised that mutations which interfere with Ub binding may also be important for recognition of SDE2 and its subsequent cleavage. To investigate this hypothesis, we prepared a panel of USP5 mutants bearing either single or double mutations, which were subsequently incubated with SDE2_FL_. Minimal differences were observed between USP5^WT^ and those mutants with intact catalytic cysteines, indicating that these mutations did not greatly impact SDE2 processing (**Figure S5C**).

We next employed ITC to compare the binding of SDE2_UBL_ to USP5 mutants. We began with a comparison between USP5^WT^ and USP5^C335A^. Mutation of the catalytic cysteine to an alanine has been previously shown to increase the affinity of select DUBs for Ub.^45^ We were therefore interested to determine if a similar change in affinity would be evident for USP5. Quantification of SDE2_UBL_ binding to USP5^WT^ (**Figure 6H**, *K*_d_=26.6 ± 5.0 µM) and USP5^C335A^ (**Figure 6A**, *K*_d_=1.67 ± 0.62 µM), revealed a striking ∼16-fold increase in affinity towards the catalytic inactive mutant, suggesting a binding mechanism analogous to that of monoubiquitin. Indeed, a similar enhancement was observed when monoubiquitin was titrated into USP5^WT^ and USP5^C335A^ (**Figure S6A/B**), consistent with reported findings.^45^ Similarly, an increase in affinity was observed by TR-FRET (**Figure S6C**). These parallels suggest that SDE2_UBL_ and Ub share a common interaction mode and likely occupy similar binding pockets. We next considered the importance of the ZnF domain. ITC measurements demonstrated a complete loss of SDE2_UBL_ binding to USP5^H232A^ (**Figure 6I**), in line with what has been reported for Ub.^46^ This supports the conclusion that, like Ub, SDE2_UBL_ relies on the structural integrity of the ZnF domain for engagement.

Despite the similarity in binding modes, we found no evidence of SDE2_UBL_-mediated activation of USP5 under any condition tested, in contrast to Ub which has been reported to stimulate USP5 catalytic activity, dependent on recognition of an unanchored C-terminal diglycine motif.^46,47^ This difference was evidenced by a fluorescence polarization cleavage assay whereby the presence of monoubiquitin increased the activity of USP5, while the addition of SDE2_UBL_ resulted in no significant difference (**Figure S7**). Thus, while SDE2_UBL_ appears to mimic Ub in its binding to USP5, it does not appear to exert the same allosteric effects, highlighting a subtle but biologically significant divergence in their functional outcomes.

## Discussion

The biological function of proteins is regulated by an intricate network of post-translational chemical and enzymatic modifications, including phosphorylation, ubiquitination, and proteolysis.^1^ Among these, proteolysis is a unique and irreversible regulatory mechanism. Historically, it has been viewed through the lens of degradation, serving to remove misfolded or damaged proteins, or to maintain cellular homeostasis *via* systems such as the ubiquitin-proteasome pathway and caspase-mediated apoptosis.^48^ However, recent advances in degradomics have fundamentally reshaped this perspective. These studies have revealed that proteolysis also plays a highly selective and instructive role in modulating protein activity, localisation and interaction across nearly all biological pathways, from signal transduction to organelle biogenesis.^49^

Within this broader regulatory framework, the nuclear protein SDE2 presents a compelling case study. Owing to its involvement in such fundamental processes, the activity of SDE2 must be stringently regulated. A key point of regulation lies in its proteolytic processing: removal of the N-terminal ubiquitin-like (UBL) domain is essential to liberate the functional C-terminal domain, which mediates the downstream biological effects of SDE2. Yet, despite the importance of this cleavage event, the identity of the responsible protease and proteolytic mechanism had remained unresolved.

In this study, we identify USP5 as the principal cysteine-dependent protease responsible for the site-specific cleavage of the SDE2_UBL_ domain. Through a combination of activity-based probe profiling, quantitative binding assays, and cell-based assays, we show that SDE2 is exclusively cleaved by USP5 both *in vitro* and in cell via a direct and specific interaction with the SDE2_UBL_ domain. This finding adds to the expanding biological repertoire of USP5. Importantly, USP5 has been implicated in homologous recombination and other DNA damage response pathways, aligning with the known roles of SDE2.^50,51^ Furthermore, genes co-expressed with USP5 in pan-cancer are known to be heavily involved in spliceosome and RNA splicing regulation, suggesting a potential complex interaction with SDE2-associated cellular pathways.^52^

Like other USP family members, USP5 isopeptidase activity facilitates the hydrolysis of the isopeptide bond between Ub chains and substrate proteins. However, as opposed to other DUBs, USP5 exhibits a unique ability to stabilise the pool of free Ub by preferentially disassembling unanchored polyubiquitin chains into monomeric Ub.^46,53^ This is specifically enabled by its unique modular domain architecture which orchestrates substrate recognition, positioning, and catalysis, allowing for the critical recognition and tight binding to substrates.^46,53^ This includes a ZnF-UBP domain that binds the free C-terminal diglycine motif of Ub and multiple UBA domains that contribute to polyubiquitin recognition. The inherent flexibility of its UBA domains, tethered to the catalytic core via disordered linkers, allows USP5 to accommodate and disassemble a broad spectrum of Ub chain types (e.g., K48-, K63-, K11-linked, and linear chains).^47,54^ Notably, while USP5 is active against polyubiquitin chains, it does not cleave the linkage between ubiquitin and substrate proteins.^55^ Therefore, its ability to recognise and cleave SDE2_FL_ highlights a dedicated and selective recognition mechanism.

Comparative assays showed that although Ub remains the preferred substrate—as demonstrated by the complete and rapid cleavage of Ub₄—SDE2 is also processed rapidly by USP5 (∼50% cleavage within 5 minutes). Additionally, its ABP (HA-SDE2_UBL_-PA) exhibits reactivity comparable to Ub-ABP and significantly higher than ISG15, another ubiquitin-like modifier involved in antiviral responses and a known non-canonical substrate of USP5.^40^ This highlights a previously unappreciated substrate promiscuity of USP5 and suggests that its substrate scope may extend beyond classical Ub derivatives under certain structural or contextual conditions.

It is also important to note the difference in the efficiency of SDE2_FL_ processing between recombinant and cellular contexts. While processing of recombinant SDE2_FL_ by USP5, and labelling of USP5 by HA-SDE2_UBL_-PA, stalled at ∼50% across multiple assays conditions, full processing and labelling were observed in cell lysate. It has been previously reported that some DUBs are activated by cofactors.^56,57^ It is therefore possible that, unlike for Ub processing, a USP5 cofactor or binding partner may be required to increase efficiency of SDE2_FL_ processing. Such a cofactor may present in cell lysate but absent in the recombinant protein context.

Using orthogonal biophysical techniques, including isothermal titration calorimetry (ITC) and TR-FRET, we quantified the interaction between USP5 and SDE2_UBL_, revealing a high-affinity binding mode reminiscent of the interaction between USP5 and monoubiquitin. Mutational analysis further demonstrated that the catalytic cysteine (C335) and the ZnF-UBP domain are critical for SDE2_UBL_ recognition, reinforcing the structural similarity to canonical Ub. Interestingly, we were unable to detect strong binding of SDE2_FL_ to USP5^C335A^. We speculate that this catalytically inactive mutant may exist in a conformationally locked state which does not accommodate the full-length protein. H232A mutation, as for Ub,^46^ reduced SDE2_UBL_ binding affinity to below the detection limits. However, the same mutation did not prevent SDE2 processing, suggesting that while an intact and flexible ZnF-UBP domain is important to accommodate a large array of possible substrates, it is not essential for catalytic activity. Therefore, a more significant structural rearrangement may be responsible for the recognition of SDE2_FL_. Additionally, we have observed that SDE2_UBL_ does not stimulate USP5 activity in the same way as monoubiquitin despite the similarities in binding, indicating that SDE2 acts as a unique pseudo-substrate.

The identification of this novel USP5-SDE2 interaction axis adds to the pool of research which highlights how the cleavage of UBLs is mediated by a suite of proteases with varied substrate specificity and domain organization. UBLs such as SUMO and NEDD8 are deconjugated by distinct families of UBL-specific proteases, including sentrin-specific proteases (SENPs), which display minimal cross-reactivity with Ub.^58,59^ Similarly, ISG15 is canonically deISGylated by USP18, a USP family member with no known activity toward Ub.^13,60^ However, more recent studies have revealed context-dependent cross-reactivity among DUBs and UBL proteases: USP16 and USP24, for instance, have been shown to deISGylate substrates in specific cellular environments.^40,61^ USP16 also cleaves Fubi, a UBL with moderate sequence similarity to both Ub and ISG15 (∼36–38%),^62^ further demonstrating that deubiquitinases can accommodate structurally related yet distinct modifiers. Our findings that USP5, a canonically ubiquitin-specific protease, proteolytically removes SDE2_UBL_ from full-length SDE2 with considerable efficiency, thus aligns with a growing body of evidence that UBL specificity is more nuanced than previously appreciated.

Alongside USP5, we also examined close paralog USP13, as a candidate for SDE2 processing. Despite sharing high sequence similarity (67.3% at the nucleotide level, 54.8% at the amino acid level), USP13 showed minimal catalytic activity toward SDE2 in both recombinant and cellular assays. HA-SDE2_UBL_-PA covalently labelled USP5 but not USP13, and knockdown of USP5, but not USP13, impaired SDE2 processing in cells. These results support a functional divergence between USP5 and USP13 and reinforce the substrate specificity of USP5 toward SDE2_UBL_. It has been reported that USP13 might exist in a constitutive self-inhibition status possibly supported by the interaction of UBA with its ZnF domain.^63,64^ It has been hypothesised that this self-inhibition status may be released by recruitment of other proteins or modification, such as phosphorylation. Therefore, we cannot exclude the possibility that under certain stimuli, USP13 might be able to process SDE2.

Collectively these findings expand the known substrate repertoire of USP5 and suggests broader implications for how ubiquitin-like proteins may be regulated by DUBs beyond their canonical roles. Further investigation into the structural determinants of this specificity, as well as the downstream consequences of SDE2 processing *in vivo*, will deepen our understanding of both USP biology and SDE2-regulated pathways.

## Methods

### Reagents used in this study

A list of plasmids used in this study can be found in Supplementary Table S1. Further information and requests for reagents should be directed to C.M. All complementary DNA (DNA) constructs in this study were generated by C.M., P.M.T., and the cloning team at the Medical Research Council Protein Phosphorylation and Ubiquitylation Unit (MRC PPU) Reagents and Services. All plasmids were deposited with the MRC PPU Reagents and Services and are available upon request at https://mrcppureagents.dundee.ac.uk/. ISG15-PA and the pTXB1_His-3C-Ub (1-75) plasmid used in this study were a kind gift from Rafael Salazar Claros of the Swatek lab.

### Cell culture

HEK293 cells, obtained from ATCC, were cultured in high-glucose Dulbecco’s modified Eagle’s medium (DMEM, 4.5 g/L glucose) supplemented with 10% fetal bovine serum (FBS), L-glutamine, and Penicillin and Streptomycin (Gibco). Cells were maintained at 37 °C in a 5% CO_2_ atmosphere. Cells were maintained for no more than 30 passages. All cell lines were routinely tested for mycoplasma contamination using MycoAlert kit from Lonza.

### Cell lysis

Cells were lysed in RIPA buffer supplemented with cOmplete EDTA free protease inhibitor cocktail (Roche). The dishes were placed on ice and the media aspirated. Tissue layer was washed twice with ice-cold phosphate saline buffer (PBS) before adding the lysis buffer. Cells were scraped from the surface and cell extract was collected. Insoluble fraction was removed by centrifugation at 4◦C (25 min, 14,000 x *rpm*). Protein concentration was measured using Pierce™ Coomassie (Bradford) Protein Assay Kit.

### SDS-PAGE and Western blotting

Samples for and SDS-PAGE and subsequent immunoblotting were heated at 95 °C for ∼ 5 min in NuPAGE LDS Sample Buffer with DTT. SDS-PAGE was performed on 4–12% Tris-Acetate NuPage® Novex® (Life Technologies) polyacrylamide gels with 3-(*N*-morpholino)propanesulfonic acid (MOPS) running buffer (ThermoFisher Scientific) at 130 V.

Samples for Western blotting were transferred to a nitrocellulose membrane using wet transfer (20% methanol, 80 V for 80 min). Membranes were blocked with 5 % w/v Bovine serum albumin (BSA) in Tris-buffered saline (TBS) with 0.1% w/v Tween-20. The following antibodies were used in this study in the given concentrations: USP5 (AB154170, abcam, 1:1000), USP13 (12577S, Cell Signaling, 1:1000), NanoLuc (N700A, Promega,1:1000), HaloTag (G921A, Promega, 1:1000), SDE2 (HPA031255, 1:1000), HA (3724S, Cell Signaling,1:1000), Flag (F7425, Sigma, 1:2000), B-actin (4970S, Cell Signaling, 1:1000), GAPDH (2118S, Cell Signaling, 1:1000), IRDye 800CW Goat anti-Rabbit Secondary Antibody (926-32211, Licor, 1:25000), IRDye® 800CW Donkey anti-Mouse Secondary Antibody (926-32212, Licor, 1:25000). Fluorescent detection was performed with a Li-Cor Odyssey CLx scanner.

### Protein expression

Competent *Escherichia coli* BL21 (DE3) cells were transformed and then used to inoculate a starter culture of TB media (200 mL) supplemented with ampicillin (final concentration 100 µg/mL) which was incubated at 37 °C with shaking (180 rpm) for ∼18 h. Expression cultures of TB (1 L, 100 µg/mL ampicillin) were inoculated with starter culture (15 mL), grown at 37 °C 180 rpm in 2.5 L high expression flasks containing 1 L of TB and 100 µg/mL ampicillin to an optical density of 0.6 at 600 nm (OD600). When the OD600 ∼ 0.6 was reached (∼ 4 h), cells were induced with 100 µM isopropyl-1-thio-β-d-galactopyranoside (IPTG) and left incubating for 24 h in the shaker at 18°C 240 rpm. Cells were pelleted by centrifuging at 4200 rpm for 30 min at 4 °C, supernatant decanted and resuspended in 50 mL (per 1 L of culture) lysis buffer (50 mM HEPES (pH 7.5), 150 mM NaCl, 20 mM imidazole, 18 µM leupectin hemisulfate, 250 µM AEBSF, 1 mM DTT), after which the pellets stored at −80 °C.

### Protein purification

Cells were lysed using an ultrasonic processor for 5 min (10 s on/ 15 s off, 60% amp) at 4°C. Supernatant was collected by centrifuging at 20 000 rpm for 30 min at 4 °C. Collected lysate was filtered through a 0.2 µm filter. From there on purification differed for different constructs.

For USP5^WT^ and mutant constructs, lysate was incubated with Ni-NTA agarose beads (5 mL per 1 L of culture lysate) for 2 h at 4°C on a roller. Loaded beads were captured in chromatography column, washed 10 CV with wash buffer A (50 mM HEPES (pH 7.5), 150 mM NaCl, 20 mM imidazole) and eluted with 10 CV of elution buffer A (50 mM HEPES (pH 7.5), 150 mM NaCl, 100 mM imidazole). Elution was concentrated down (Amicon^®^ Ultra Centrifugal Filter) to injection volume (<5 mL) for a size exclusion chromatography (SEC) by gel filtration on a HiLoad 16/600 Superdex 200 pg column (Cytiva) using gel filtration buffer (50 mM HEPES (pH 7.5), 150 mM NaCl, 5% glycerol, 1 mM DTT). Fractions containing the USP5 were pooled and concentrated by ultrafiltration.

For constructs containing SUMO [His-SUMO-SDE2_FL_(1-451), His-SUMO-SDE2_CT_(78-451) and His-SUMO-SDE2_UBL_(1-77)], protein was captured with Ni-NTA agarose beads (2 h, 4°C), washed in chromatography column with wash buffer A (10 CV), eluted with elution buffer A (10 CV) and set up for overnight dialysis into no imidazole buffer (50 mM HEPES (pH 7.5), 150 mM NaCl, 1 mM DTT) with His-tagged SUMO-protease (1 µg protease per 2 mg of protein). Dialysed solution was passed through fresh Ni-NTA agarose beads to capture SUMO-protease and uncleaved protein. Flowthrough (His-SUMO tag cleaved protein of interest) was concentrated down to injection volume for gel filtration. SDE2_FL_ and SDE2_CT_ were run on HiLoad 16/600 Superdex 200 pg column (Cytiva), while SDE2_UBL_ was run on HiLoad 16/600 Superdex p75 column (Cytiva) (using gel filtration buffer). Fractions containing the desired construct were pooled, concentrated, flash-frozen, and stored at −80°C.

HA-SDE2_UBL_(1-77), lysate was incubated with chitin agarose beads (New England) for 3 h at 4°C on a roller. Loaded resin was washed with 10 CV of wash buffer B (50 mM HEPES (pH 7.5), 150 mM NaCl) and then incubated in a cleavage buffer (50 mM HEPES (pH 7.5), 150 mM NaCl, 50 mM DTT) over 72 h at 4°C. The elution was collected in a falcon tube along with a 1 CV resin wash (wash buffer B). Elution was concentrated and gel filtrated with a HiLoad 16/600 Supedex p75 column (Cytiva). Fractions containing POI were pooled, concentrated, flash-frozen, and stored at −80°C.

SUMO protease (used for the beforementioned His-SUMO tagged SDE2 constructs) was expressed and purified by Ni-NTA affinity a previously described.^65^ In short, lysate was incubated with Ni-NTA agarose beads for 2 h at 4°C, captured on affinity column, washed 10 CV with wash buffer A, eluted 10 CV with elution buffer A, concentrated down, and stored in −80°C.

### Lysate fractionation and screening

*Lysis, Heparin and SourceQ fractionation:* Expi293 cells were grown in 100 mL suspension in A14351-01 media were grown until a confluency of 5×10^6^ cell/mL. Cells were washed in PBS and resuspended in 5 volumes of hypotonic buffer (50 mM Tris pH 7.5, 7 mM-2-BME, 1 mM TCEP, 1 mM AEBSF, 10 µg/mL Leupeptin), and incubated on ice for 20 min before lysis by mechanical stress through a douncer homogenizer followed by (>20 sequential passes through 21-23-gauge needles). Resulting lysate was cleared by centrifugation (20 min, 13,000 rpm) and soluble fraction was filtered through 1 Whatman syringe filter (PVDF 0.45 µM). The lysate was buffer exchanged to MOPS-LSB (30 mM MOPS pH 7.0, 10% glycerol, 7 mM 2-ME, 1 mM TCEP, 0.015% Brij35) using a 20 mL Sephadex G25 fine column (XK16/10) using an Akta Pure Fast Protein Liquid Chromatography (FPLC) device. The lysate was applied to a 1 mL Heparin HiTrap-HP column and proteins were eluted with an increased gradient of a high salt buffer (30 mM MOPS pH 7.0, 1.2 M NaCl, 5% glycerol, 1 mM DTT, 0.015% Brij 35) and collected as 1 mL fractions. The pH of the flowthrough from the Heparin chromatography was adjusted to pH 8.0, by addition of 300 µL (1% v/v) of unbuffered 1 M Tris-Base and it was passed through a 1 mL Source 15 Q column (HR10/10) and eluted on a salt gradient (30 mM Tris pH 8.0, 10% glycerol, 7 mM 2-ME, 1 mM TCEP, 0.015% Brij35, 1 M NaCl). SourceQ fractions (1mL volume) were eluted into a 96 well block at 1 mL intervals. Heparin and SourceQ binding fractions were immediately snap-frozen in liquid nitrogen and stored at −80 °C until use.

*Fraction screening against SDE2_FL_:* Lysate, fractions from Heparin and SourceQ, and flowthrough (3ml) were diluted with 10ml of assay buffer (50 mM Tris pH 7.5, 1 mM DTT) and treated with either 2 mL of a solution (22.8 mg/ml) of recombinant SDE2_FL_ or 2 mL of buffer. Samples were incubated at 30 °C for 30 min before quenching with 4× SDS loading dye. Samples were boiled at 95 °C for 5 min and resolved by SDS-PAGE on 4–12% Tris-Acetate NuPage® Novex® (Life Technologies) polyacrylamide gels with 3-(*N*-morpholino)propanesulfonic acid (MOPS) running buffer (ThermoFisher Scientific) at 130 V. Gels were stained with Comassie Instant Blues before imaging on ChemiDoc (Biorad).

*Chromatography over Superdex 200:* The active fractions A12 and B1 were found to contain the relevant enzyme activity. Fractions were concentrated to 0.4 – 0.5 mL and individually purified on a Superdex 200 Increase GL 300 10 column equilibrated in 30 mM Tris pH 7.5, 10% glycerol, 150 mM NaCl, 7 mM 2-BME, 1 mM TCEP, 0.015% Brij35. Fractions (1 mL) were collected and subjected to screening as described above. Fractions showing processing of SDE2_FL_ (B2, B3 and B4) together with fractions A11, A12, B1 and B5, B6, B7 were processed for mass spectrometry analysis.

### High-throughput MALDI-TOF/MS SDE2 cleavage assay

A panel of recombinantly expressed DUBs were tested for their ability to cleave SDE2_UBL_ from SDE2_FL_. The screening was performed and analysed similarly to previously reported.^66^ Briefly, recombinantly expressed DUBs were diluted in 40 mM Tris-HCl pH 7.5, 0.01% Bovine Serum Albumin (BSA) and 1 mM TCEP to a final concentration of 50-200 ng/µL. The reaction was started by adding either positive control substrates (Ub dimers, trimers of different linkages or other pseudo-substrates as previously reported^66,67^) at the final concentration of 1.2 - 2.4 µM or SDE2_FL_ at final concentration of 3 µM. The reaction was stopped after 30- and 60-minutes incubation time using a mixture of 2% TFA and 0.8 µM ^15^N Ub (as internal standard). Positive control activity was reported as ratio of Ub/^15^N Ub (integrated areas under the peak of 8565.76 and 8669.47 *m/z*) while activity toward SDE2_UBL_ was assessed as ratio of SDE2_UBL_/^15^N Ub (SDE2_UBL_ 8348.9 observed *m/z*). Samples were mixed 1:1 with 2′,6′-Dihydroxyacetophenone matrix (DHAP), spotted on a 1536 AnchorChip MALDI-target using a Mosquito nanoliter pipetting system and analysed using a rapifleX MALDI Tissuetyper MS System as previously reported.^66^

### Alanine scanning

Human embryonic kidney 293T (HEK293T) cells were seeded in complete growth medium (2ml of DMEM + 10% Fetal Bovine Serum, 2mM L-glutamine, 2% Pen/strep) into 6 well plates at a density of 0.4 x 10^6^ cells/well to achieve 70% density on day of transfection. 155 µL of 0.020 µg/µL plasmid solution was prepared in OptiMEM® in an Eppendorf tube for each plasmid, and 9.8 µL of FuGENE® HD reagent added. Tubes mixed carefully and incubated for 15 minutes at room temperature. 150 µL of each transfection complex was added to individual wells of the seeded plates, before incubation for 48 hours at 37 °C.

### Preparation of activity-based probes

HA-SDE2_UBL_-PA was generated through C-terminal functionalisation via an intein approach as previously described for other UBLs. An activated C-terminal MESNA thioester, HA-SDE2_UBL_-MESNA, was utilised as a key intermediate.

An N-terminally tagged HA-tagged human SDE2(1-76) insert was cloned into a pTXB1 vector using NedI and SapI restriction enzymes to generate pTXB1_SDE2(1-76). Protein expression was carried out according to the general procedure described above. To generate HA-SDE2_UBL_-MESNA, cells pellets were resuspended in lysis buffer (50 mM HEPES, 150 mM NaCl, 0.5 mM TCEP, pH 7.5) and sonicated on ice. Lysate was clarified by centrifugation (20,000 rpm, 30 min at 4 °C), followed by filtration (0.22 µm syringe filter). Clarified lysate was incubated with chitin beads (NEB #S6651) for 4 h at 4 °C on a rotatory platform. After incubation, beads were collected in a reusable column and washed with lysis buffer (10× column volume). Three column volumes of cleavage buffer (50 mM HEPES, 150 mM NaCl, 0.5 mM TCEP, 100 mM MESNA, pH 7.0) was added to the beads, followed by incubation for ∼62 h at 4 °C on a rotatory platform. The protein was then eluted from the column and concentrated using a centrifugal filter (Amicon® Ultra, 15 mL, 3 kDa MWCO). The HA-SDE2_UBL_-MESNA product was then either purified by RP-HPLC (for long-term storage at −80 °C or directly conjugated with propargylamine.

To generate HA-SDE2_UBL_-PA, a solution of HA-SDE2_UBL_-MESNA was treated with propargylamine (final concentration of 225 mM). The solution was incubated at 25 °C for 2 h and reaction completion monitored by intact LCMS. The final product was isolated by RP-HPLC and lyophilised to give a white powder (∼2.5 mg/L of culture). The ABP was renatured by dissolving the powder in minimal DMSO, followed by the slow addition of buffer (50 mM HEPES, 150 mM NaCl, 0.5 mM TCEP) to a final DMSO concentration of 5%. Using a centrifugal filter (Amicon Ultra, 0.5 mL, 3 kDa MWCO), the refolded probe was concentrated, diluted with buffer, and concentrated again to remove DMSO. Final concentration was determined using A280 absorbance (NanoDrop One, ε = 7365 M^−1^cm^−1^)

His-Ub-PA was generated as described for HA-SDE2UBL-PA.

### RP-HPLC

Reverse-phase HPLC purification was performed on a ThermoFisher UltiMate 3000 HPLC system coupled to a Phenomex Jupiter 5 µm C4 300 °A LC column (250 x 10 mm). Proteins were concentrated to a maximum injection volume of 3.5 mL and subject to a gradient of 10-80% MeCN (0.1% TFA) in H_2_O (0.1% TFA) over 30 min with absorbance was monitored at 214 and 280 nm. Data was analysed using Chromeleon 6.80. Intact LCMS was used to identify fractions containing the desired product, which were pooled and lyophilized.

### Intact LCMS

Intact LCMS was performed using a Zorbax 300SB-C3 5 µm (150 × 2.1 mm) column coupled to an Agilent system composed of a 1260 Bin Pump VL and Degasser, a 1290 Thermostatted Column Compartment and a 6130 Quadropole LC/MS. Sample injection volumes ranges between 1-5 µL and subject to a gradient of 10-75% MeCN (0.05% TFA) in H_2_O (0.05% TFA) over 20 min. Data analysis, including deconvolution of mass ions, was performed using Agilent LCMSD ChemStation (Rev. B.04.03-SP1)

### SDE2 cleavage and ABP labelling assays

Both SDE2 cleavage and ABP labelling assays were performed in 0.5 or 2 mL Eppendorf tubes by diluting purified DUBs (1 µM final concentration) and ABPs or SDE2_FL_ (2 µM final concentration) in reaction buffer (50 mM Tris, 1 mM DTT, pH 7.5). The reactions were incubated at 37 °C for the indicated time then quenched with 4× LDS/DTT and analysed by SDS-PAGE.

### Activity-based DUB profiling

*Cell lysis and probe labeling:* HEK293 cells were cultured in 15 cm dishes until ∼70-80% confluent. The cells were collected by rinsing the plate with ice-cold PBS, collected into a centrifuge tube and pelleted by centrifugation at 4 °C (3 min, 350 × g). The resulting pellet was washed twice with PSB before being flash-frozen in liquid nitrogen and stored at −80 °C until lysis. Frozen cells were resuspended in lysis buffer (50 mM Tris, pH 7.5, 5 mM MgCl_2_, 0.5 mM EDTA, 250 mM Sucrose, 1 mM DTT). One volume of acid-washed glass beads (Sigma Aldrich, G4649) per 3 volumes of ice-cold glass bead buffer was added to the tube containing the resuspended cells followed by 10 vortexing/cooling cycles (30 s bursts followed by incubation on ice for 2 min). The lysate was cleared by centrifugation (14,000 g, 4 °C, 25 min). The protein concentration was determined by Bradford assay. Cell extract (500 µg) was diluted to a final volume of 300 µL using lysis buffer and incubated with 1 µM of probe for 45 min at 37 °C and 300 rpm. The reaction was quenched by adding 0.4% SDS and 0.5% NP-40. To immunoprecipitated DUB–ABP complexes, 50 µL of anti-HA magnetic beads slurry (Thermo Scientific), previously washed four times with NP-40 lysis buffer, was added to 105 µL samples diluted with 210 µL of NP-40 lysis buffer and incubated on a rotator overnight at 4◦C. Beads were pelleted by placing the sample on a magnetic rack, and the supernatant was removed. Beads were washed four times with 300 µL of NP-40 lysis buffer. Protein complexes were eluted by boiling beads in 80 µL of 2× LDS sample buffer at 75 °C for 15 min and 10% were analysed by Western blotting. 25 µL of the DUB-ABP immunoprecipitated samples were processed for proteomic analysis as described below.

### Mass spectrometry proteomics experiments

*Sample preparation*: Samples (25 µL) were reduced with 5 mM Bond-Breaker TCEP solution for 15 min at 55 °C. Samples were alkylated with 20 mM iodoacetamide for 10 min at room temperature, followed by acidification (final concentration 2.5% phosphoric acid). Sample were diluted by adding 165 mL of binding buffer (100 mM TEAB in 90% methanol). Samples were applied to S-trap micro column (ProtiFi) and spun down at 4,000 × g for 30 sec to trap proteins. Spin columns were washed 3 times with 150 mL of binding buffer and centrifuged in between each wash (4,000 × g for 30 s) followed by a last centrifugation for 1 min to dry the spin column membrane from residual washing buffer. Digestion buffer (20 mL, 50 mM TEAB) containing trypsin (5 mg) was added to the top of the S-Trap column and incubated overnight at 37 °C. Digested peptides were eluted in three subsequent steps using buffer 1 (50 mM TEAB in water), buffer 2 (0.2% formic acid in water), and buffer 3 (50% acetonitrile in water). Eluted peptides were combined and dried down in a SpeedVac.

*LC-MS Analysis:* Samples were dissolved in HPLC-grade water containing 0.1% formic acid and 0.015% n-dodecyl β-D-maltoside. Peptide separation was performed on a Vanquish Neo UHPLC system in trap-and-elute mode, interfaced with an Orbitrap Astral Mass Spectrometer (Thermo Fisher Scientific). Peptides were initially captured on a PepMap Neo Trap Cartridge (Thermo Fisher Scientific, #174500) and subsequently separated using a C18 EASY-Spray HPLC Column (Thermo Fisher Scientific, #ES906). The elution gradient spanned 11.8 minutes, increasing from 1% to 55% buffer B (Buffer A: 0.1% formic acid in water; Buffer B: 0.08% formic acid in 80:20 acetonitrile:water) with the following profile: 0.7 min at 1.8 µL/min from 1% to 4% B, 0.3 min at 1.8 µL/min from 4% to 8% B, 6.7 min at 1.8 µL/min from 8% to 22.5% B, 3.7 min at 1.8 µL/min from 22.5% to 35% B, and a final step of 0.4 min at 2.5 µL/min from 35% to 55% B. Peptides were analysed via data-independent acquisition (DIA) on the mass spectrometer.

*Data Processing and Statistical Analysis:* Peptide identification was carried out against the UniProt Swiss-Prot human database (release date: 02/01/2023) using DiaNN (v1.9)^68^ in library-free mode, with cross-run normalization disabled. Data analysis was performed using Python (packages: pandas, numpy, sklearn, scipy, rpy2, Plotnine, and Plotly) and R (Limma package^69^). Protein groups were retained only if at least two peptides were identified and quantified in a minimum of two replicates. Missing values were imputed using a Gaussian distribution centred on the median, applying a downshift of 1.8 and a width of 0.3 (relative to the standard deviation). Differential protein regulation was assessed using LIMMA eBayes, with multiple hypothesis correction via the Benjamini-Hochberg method. Proteins were considered significantly regulated if the adjusted p-value was below 0.05 and the fold change exceeded 1.5 or was lower than 1/1.5. The mass spectrometry proteomics data have been deposited to the ProteomeXchange Consortium via the PRIDE^70^ partner repository with the dataset identifiers PXD064153 and PXD064155.

### Endogenous siRNA

U2OS cells were seeded at a density of 0.4 x 10 6 cells/well into 6-well plates to achieve 70% density on the day of transfection. USP5 #2 (cat. D-006095-02-0005, Dharmacon), USP5 #3 (cat. D-006095-03-0005, Dharmacon), USP5 pull (cat. L-006095-00-0005, Dharmacon), USP13 pull (cat. D-006064-00-0005, Dharmacon), SDE2 (cat. L-016456-02-0005, Dharmacon) and control siRNA (cat. D-001820-10-05, Dharmacon) were used. SiRNA (4µg) were diluted in OptiMEM and mixed with a solution of 20µl of X-tremeGENE siRNA transfection reagent and 80µl of OptiMEM. Solutions were mixed and incubated at room temperature for 20 min. Transfection reagent:siRNA complex was added dropwise to the cells and the plate was swirled to ensure distribution over the entire plate. Cells were incubated for 48h before harvesting.

### nanoBRET assay

Wild-type full length SDE2 and uncleavable mutant (G76A G77A) were cloned into pF-NH Flexi Vector (Promega) with an N-terminal Nanoluc and C-terminal Halotag. HEK293T cells were seeded into 6 well plates in complete growth medium (2ml of DMEM + 10% Fetal Bovine Serum, 2mM L-glutamine, 2% Pen/strep) at a density of 0.7 x 10^6^ cells/well to achieve 70% density on day of transfection. USP5 (cat.) USP13 (cat.) and control siRNA (cat.) were used. All siRNA (5 µL of a 10µM solution) were diluted in 250 µl of OptiMEM no phenol red, 20µL RNAiMAX® (20 µL) was mixed with OptiMEM® (no phenol red) (1 mL) and 255µL dispensed into each prediluted siRNA. Tubes were mixed, incubated at room temperature for 20 minutes, then briefly centrifuged. 255 µL of each complex was added in duplicate to 6 well plate wells and incubated at 37 °C for 36 hours. 2 µg of each plasmid solution or NanoBRET™ Positive Control Vector were prepared in 100 µL OptiMEM® Reduced Serum Medium, no phenol red, in Eppendorf tubes. 6 µL of FuGENE® HD Transfection reagent was added, tubes mixed carefully and incubated for 10 minutes at room temperature. 150µL of each transfection complex was added to individual wells, before incubation for 20 hours at 37 °C, 5% CO_2_. Each well of cells washed twice in 1ml DPBS before trypsinization in 0.5 mL of 0.05% trypsin-EDTA and incubation at room temperature until cells lifted. 2 mL cell culture medium was added, and cells were collected and transferred to 15 mL conical tubes. Cells were spun down at 125 × g for 5 minutes, cell culture medium discarded and cell pellets resuspended in an equal volume of assay medium (OptiMEM® Reduced Serum Medium, no phenol red + 4% FBS). Cell counts were taken, and densities adjusted to 2 × 10^5^ cells/ml in assay medium. Each conical tube of cells was divided into four pools of 500µL and HaloTag® NanoBRET™ 618 Ligand or DMSO vehicle was added as follows: 0.5 μL ligand in two pools, 0.5 μL DMSO in one pool, and one pool left untreated. This allowed conditions to be tested in triplicate (100 μL × 3 + 2 extra) and excess cells remaining in conical tubes were washed and lysed for western blotting to verify protein expression. 100 µL of each pool of cells was plated into 3 wells of a 96 well plate. A 5× solution of NanoBRET™ Nano-Glo® Substrate was prepared in Opti-MEM® Reduced Serum Medium, no phenol red. For 60 wells, 1.75 mL Opti-MEM® Reduced Serum Medium, no phenol red, containing 17.5 μL NanoBRET™ Nano-Glo® Substrate was prepared. 25 µL of substrate was added to cells using a multidispenser. The plate was inserted into a Promega GloMax® plate reader and protocol BRET: NanoBRET 618 protocol was run with prior shaking for 30 sec. Raw NanoBRET™ units were converted into milliBRET units (mBU) by multiplying by 1000. The mean NanoBRET™ ratio was then determined and the mean corrected mBU was calculated as follows: **Mean mBU experimental – Mean mBU no-ligand control = <u>Mean corrected mBU.</u>** All graphs created using GraphPad Prism. Standard error of the mean calculated and displayed as error bars, where appropriate.

### Thermal shift assay

Components used in thermal shift assays were prepared in reaction buffer (PBS, 1 mM DTT) and the assays performed in a white BioRad PCR plate, sealed with optically clear plate sealer (BioRad). USP5 (6 µL, 4 µM) was mixed with ABP (SDE2_UBL_-PA, ISG15-PA, Ub-PA, 6 µL, 8 µM), or buffer and incubated at 37 °C for 1 h. To the protein solution was added SYPRO orange (8×, 12 µL) and plate thermocycled using a Bio-Rad CFX Opus 384 RT-PCR (10-95 °C, 0.5 °C increase every 30 s) and the fluorescence detected using the FRET channel. RFU was plotted against temperature and the first derivative (dRFU/dT) was calculated using GraphPad Prism 9. Melting temperature (T_m_) is calculated as the temperature which gives the maximum dRFU/dT value. Experiments were performed in triplicate and replicates used to calculate a mean T_m_.

### ITC

Experiments were performed on an ITC200 micro-calorimeter (GE Healthcare) at 298 K. To prepare samples, protein stock solutions were dialysed overnight in ITC buffer (2 L, 50 mM HEPES, 100 mM NaCl, 1 mM TCEP, pH 7.5) using D-Tube Dialyzers (Merck, MWCO 3.5 kDa or 11-12 kDa). The dialysis buffer was filtered and degassed then used to dilute proteins to the indicated concentration. Titrations were performed with a single injection of 0.4 µL (data point discarded) followed by 19 injections of 2 µL at 120 s intervals. Data was processed using the MicroCal ORIGIN software package. Experiments were performed in duplicate.

### TR-FRET

TR-FRET assay buffer (20 mM HEPES, 150 mM NaCl, 1 mM TCEP, 0.5% DMSO and 0.05% Tween-20, pH 7.5) was used to dilute all samples to 4 × final concentration. To each well of a 384-well white OptiPlate (Perkin Elmer) was added His-tagged donor protein (4 µL), LANCE Eu-W1024 Anti-His_6_ antibody (4 µL, final concentration 2 nM, Revvity AD0401), Cy-5 labelled acceptor protein (4 µL), and additional TR-FRET assay buffer (4 µL). Following 1 h incubation at room temperature the FRET ratio was measured at λ_exc_ 337 nm, λ_em_ = 620 nm and 665 nm using a PHERAstar FXS plate reader (BMG Labtech). All experiments were performed in triplicate. The emission values were converted to a ratio of the emission at 665 nm to 620 nm (FRET ratio = [665 nm/620 nm]*10000) and a mean value for each condition was calculated. A Cy5 labelled acceptor only control was used to determine background signal and subtracted from final readings. The one site-specific binding equation was used on Prism 10.3.1 to calculate the B_max_ and *K*_d_ for each condition.

### Fluorescence polarisation (FP)

FP assays were performed in 384-well black plates (Greiner) using FP assay buffer (50 mM HEPES, 150 mM NaCl, 1 mM DTT). Stock USP5^WT^, Ub and SDE2_UBL_ were diluted in assay buffer to 1.5 µM (3 × final concentration) and Ub-KG-TAMRA (UbiQ, UbiQ-012) was diluted to 1 µM (10 × final concentration. To assay buffer (4 µL) was added 3× USP5^WT^ (5 µL) and either 5 µL of 3× Ub, 3× SDE2_UBL_, or buffer. A blank (buffer only) and a negative control (14 µL buffer + 1 µL 10× Ub-KG-TAMRA) were also prepared. The reaction was initiated by the addition of 10× Ub-KG-TAMRA (1 µL) to each of the test wells. Using a ClarioStarPlus plate reader, fluorescence polarisation was measured at ambient temperate every 20 s for 60 min (Ex: 540, Em: 590, target mP= 150, blank corrected). Experiments were performed in triplicate. Data was analysed and visualised on Prism 10.3.1 and final polarisation values were compared using a one-way ANOVA test.

## Acknowledgements

This study was funded by the Biotechnology and Biological Sciences Research Council (BBSRC Discovery Fellowship BB/T009454/1 to C.M.), The Royal Society (Research Grant RGS\R2\212269 and University Research Fellowship URF\R1\231436, both to C.M.) and UK Medical Research Council ([MC_UU_00038/1] to Prof. Dario Alessi). P.M.T. was supported by a PhD studentship from the UK Medical Research Council (MRC) (MC_ST_00038). O.C. was supported by a PhD studentship from EPSRC MoSMed CDT and Bristol-Myers-Squibb (EP/S022791/1). We thank Rafael Salazar Claros and Dr Kirby Swatek for sharing the ISG15 probe and Ub-encoding plasmids, Dr. Pawel Lis and Prof. Dario Alessi for providing Expi293 cell pellet and support staff at MRC-PPU reagents and services. We are also grateful to Prof. Satpal Virdee for access to LC-MS and HPLC instrumentation, and to the super users of the Protein Purification Facility at the Centre for Targeted Protein Degradation for access to the ÄKTA purification system. We thank Dr. Mathieu Soetens for valuable discussions. Figures were created in BioRender (https://BioRender.com)

## Author contributions

**Conceptualization, Funding acquisition, Project administration, Supervision:** C.M.; **Methodology:** C.M., L.T.H., P.M.T., R.T., V.D.C., A.K., F.L.; **Validation:** C.M., L.T.H., P.M.T.; **Formal Analysis:** C.M., L.T.H., P.M.T., A.W., R.B., A.S., F.L.,V.D.C.; **Investigation:** C.M., L.T.H., P.M.T., A.W., R.B., A.S., A.K., R.T., F.L., M.P., V.D.C., O.C.; **Resources**: C.M., L.T.H., P.M.T., A.W., V.D.C., R.T., O.C.; **Data Curation:** F.L.; **Writing – original draft:** C.M., L.T.H., P.M.T.; **Writing – review and editing:** all authors. **Visualization:** L.T.H, P.M.T, C.M.

