## Supplementary Information for "A Proteolytic Switch: USP5 controls SDE2 function via UBL-directed cleavage"

Liam T. Hales[1*], Paul M. Tammiste[1*], Adam J. Walker[1], Rebecca Bryce[1], Andrew Sugden[1], Axel Knebel[1], Rachel Toth[1], Fred Lamoliatte[1], Marian Péteri[1], Oliwia Curry[2], Virginia De Cesare[1], Chiara Maniaci[1,2✉]

[1]MRC Protein Phosphorylation and Ubiquitylation Unit, Sir James Black Centre, School of Life Sciences, University of Dundee, Dundee, DD1 5EH United Kingdom

[2]School of Natural and Environmental Sciences, Newcastle University, Newcastle Upon Tyne, NE1 7RU, United Kingdom.

* These authors contributed equally: Liam T. Hales, Paul M. Tammiste

### Supplementary Figures

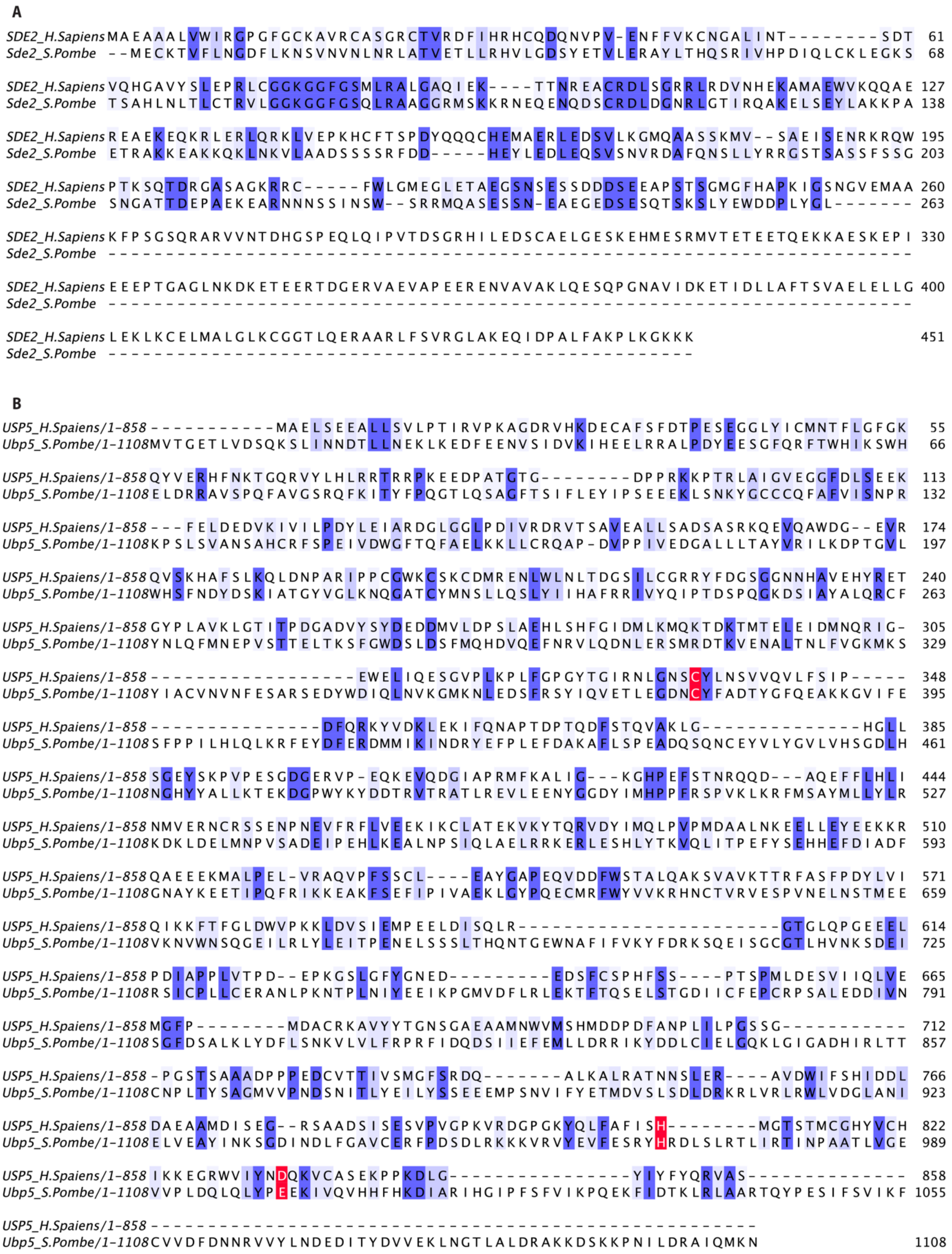


**Figure S1 | Sequence alignment of (A) *H. sapiens* SDE2 with *S. pombe* Sde2 and (B) *H. sapiens* USP5 with *S. pombe* Ubp5.** Sequences were aligned with Clustal Omega[1] and conserved residues were annotated in Jalview[2] by conservatism. Dark blue indicates fully conserved, while light blue shows semi-conserved residues. Protease catalytic triad is highlighted in red.

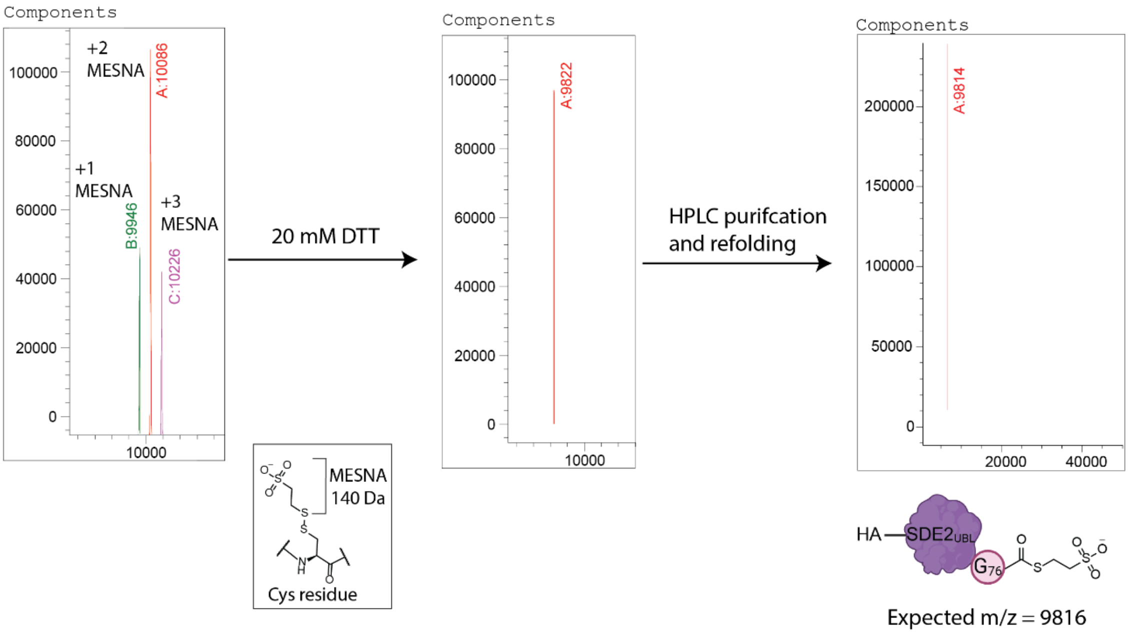


- Difference of m/z=140 between peaks
- Potentially mono-, di- and tri-MESNA reduced cysteines
- Removed via reduction with DTT (C-terminal thioester unaffected)

**Figure S2 | Intact LCMS of HA-SDE2$_{UBL}$-MESNA following the MESNA-mediated cleavage of HA-SDE2$_{UBL}$ΔG77 from the chitin resin resulted in mass ions higher than expected.** The additional masses increased by 140, which corresponds to the mass of one (140 Da) or multiple MESNA molecules. Addition of DTT lead to the disappearance of these mass ions, suggesting that they are due to the formation of disulfide bonds between Cys residues and MESNA which are reduced in the presence of DTT. HPLC purification and protein refolding then intact LCMS analysis gave a *m/z* signal corresponding to the oxidised HA-SDE2$_{UBL}$-MESNA molecular weight.

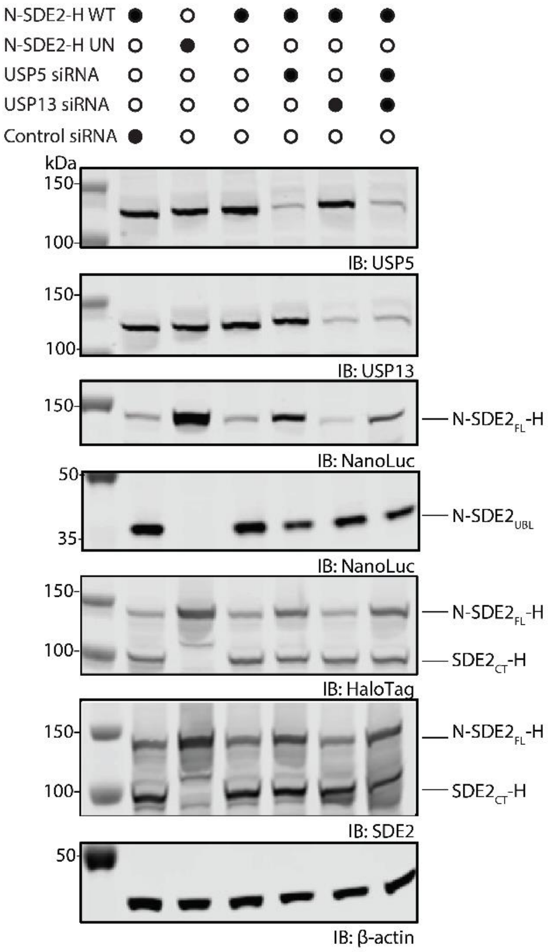


**Figure S3 | Western blot analysis of cells used for NanoBRET®.** Cells overexpressing NanoLuc-SDE2$_{FL}$-HaloTag (N-SDE2-H WT) or G76A G77A uncleavable mutant (N-SDE2-H UN) treated with USP5 or USP13 specific siRNA, singularly or in combination. Control siRNA was used to assess potential non-specific effects.

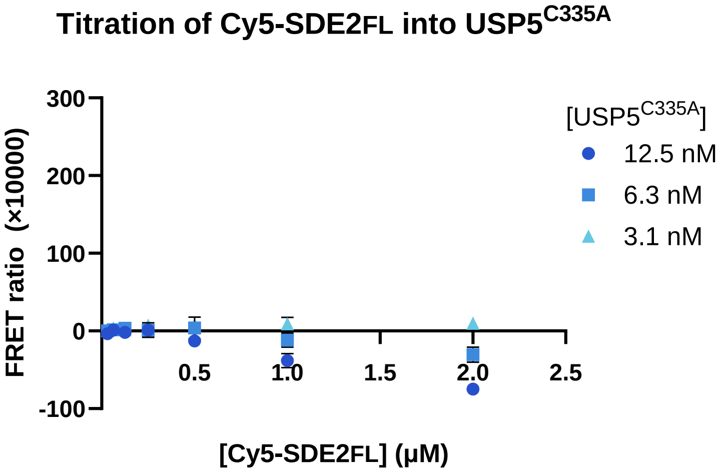


**Figure S4 | TR-FRET assessment of SDE2$_{FL}$.** Cy5-SDE2$_{FL}$ was titrated into USP5$^{C335A}$ (at 3.1, 6.3 or 12.5 nM) and the FRET ratio was measured. TR-FRET experiments were performed as technical triplicates.

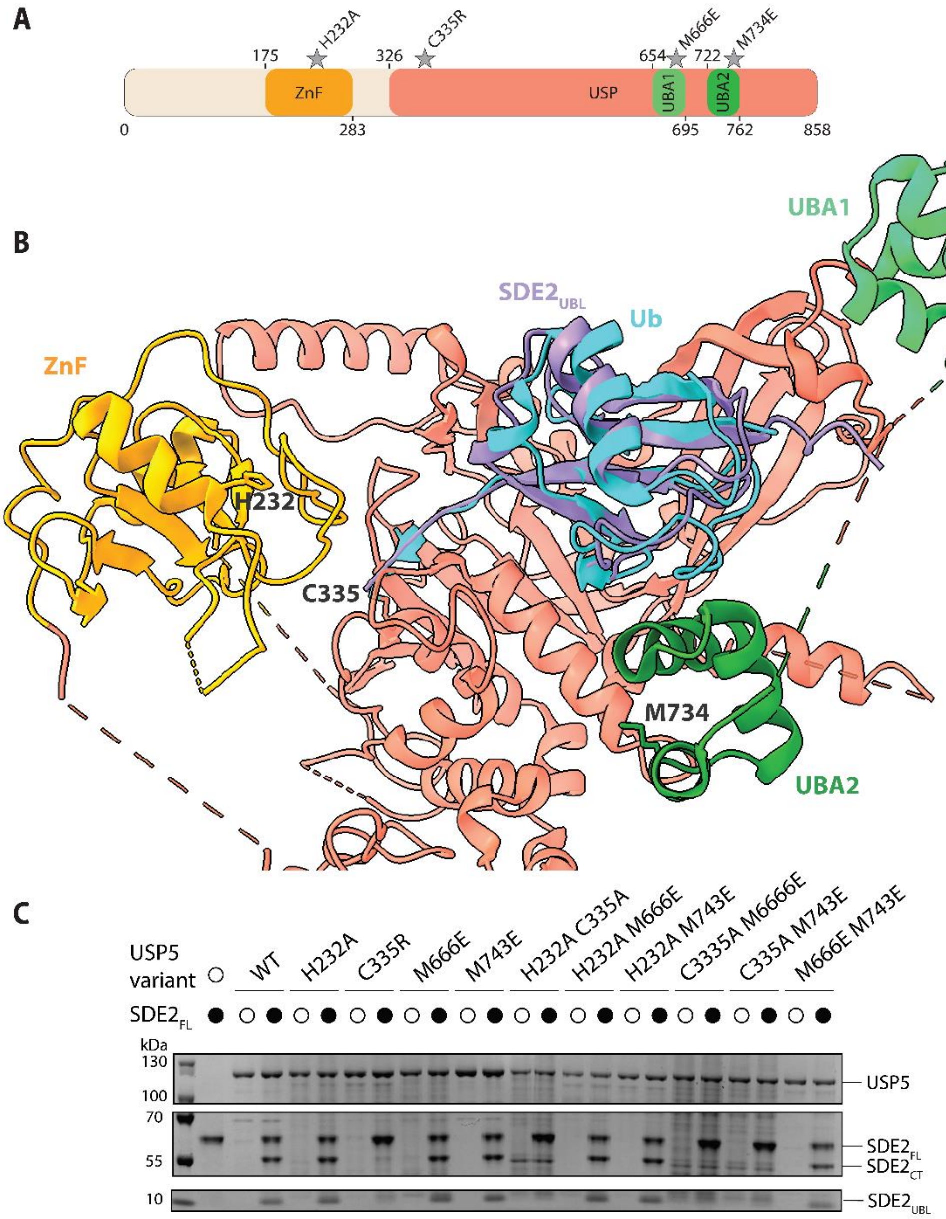


**Figure S5 | Mutation of USP5 residues important for recognition of Ub do not significantly hinder SDE2 cleavage. (A)** Schematic representation of USP5 domains their mutations and location. **(B)** $SDE2_{UBL}$ was modelled in a USP5 bound state using AlphaFold 3.[3] $SDE2_{UBL}$ structure was extracted and superimposed onto Ub (RMSD = 2.029) in a solved USP5-Ub crystal structure (PDB: 3IHP). Legend: Ub – cyan blue, $SDE2_{UBL}$ - purple, USP5 – light tomato red, ZnF – orange, UBA1 – light green, UBA2 – dark green. Mutations were introduced in each of the key Ub binding sites and are indicated. **(C)** A panel of USP5 mutants (1 µM) bearing single and double mutations were incubated with $SDE2_{FL}$ (2 µM) for 15 min at 37 °C. Cleavage of $SDE2_{FL}$ was visualised by SDS-PAGE and Coomassie staining.

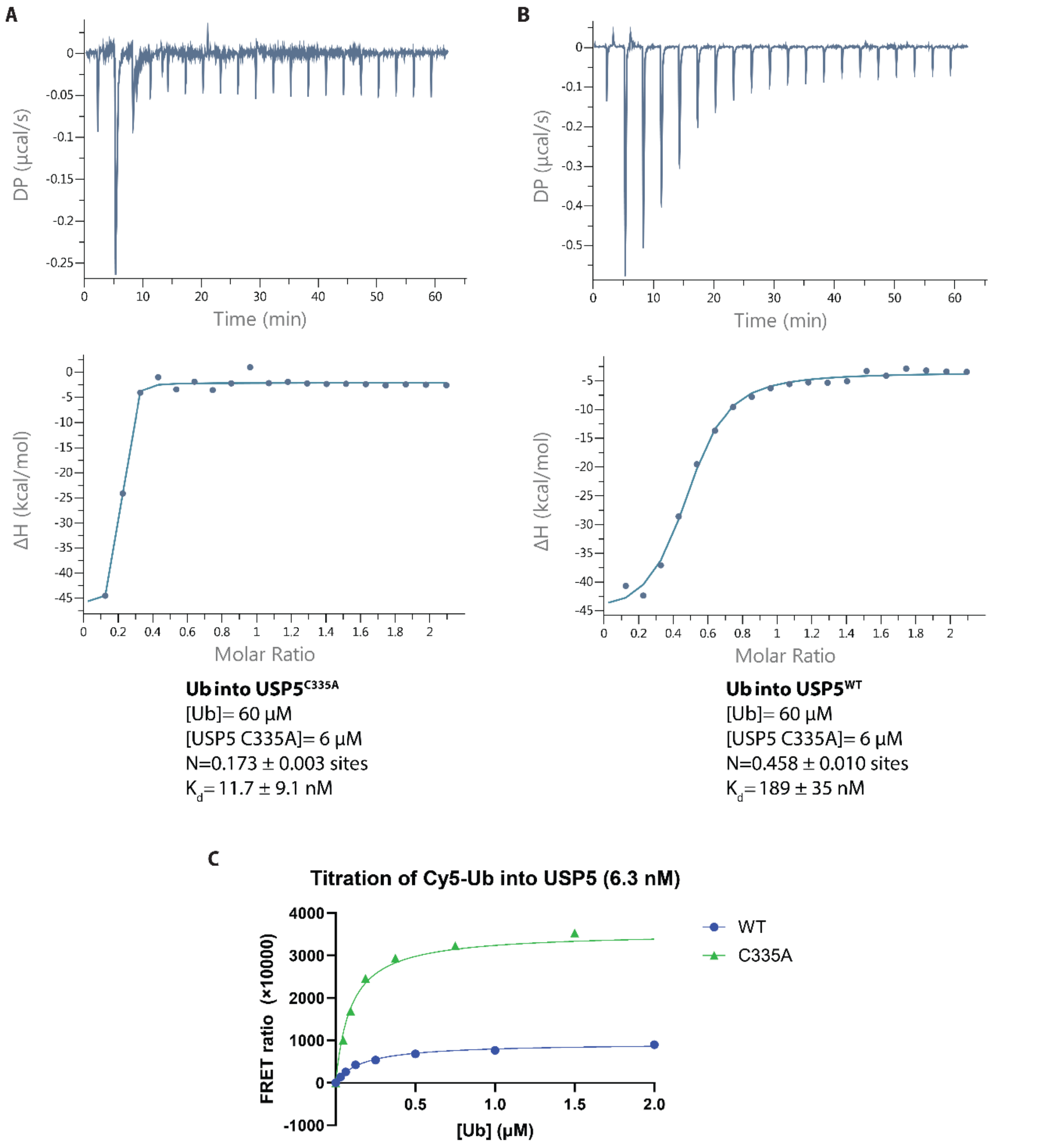


**Figure S6 | Assessment of the binding interaction between monoubiquitin (Ub) and $USP5^{WT}$ and C335A by ITC and TR-FRET. (A-B)** Using ITC, Ub (60 µM) was titrated into $USP5^{WT}$ (60 µM, A) or $USP5^{C335A}$ (60 µM, B). **(C)** TR-FRET assessment shows Ub binds to $USP5^{C335A}$ ($K_d$= 95 ± 20 nM) with ~2-fold higher affinity than $USP5^{WT}$ ($K_d$= 171 ± 20 nM). Ub was titrated into $USP5^{WT}$ (6.3 nM) or $USP5^{C335A}$ and the FRET ratio of 665/620 nm was detected. TR-FRET experiments were performed as technical triplicates and curves fitted using a one-site specific curve to calculate a $K_d$.

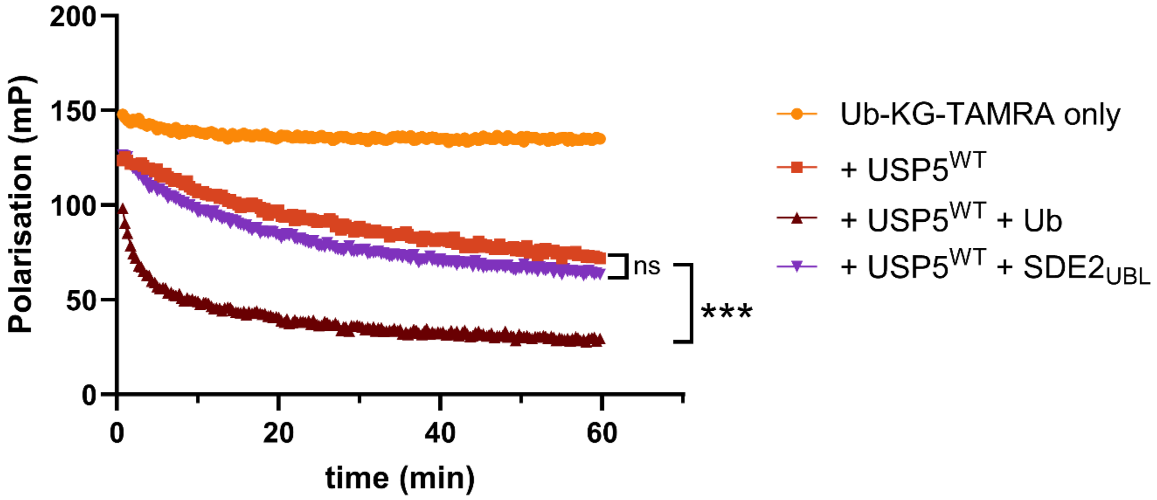


**Figure S7 | Fluorescence polarisation cleavage assay demonstrates that Ub, but not SDE2$_{UBL}$, activates USP5$^{WT}$.** Ub-KG-TAMRA (100 nM) was incubated with USP5$^{WT}$ (500 nM) for 60 min at 37 °C, either alone or in the presence of Ub (500 nM) or SDE2$_{UBL}$ (500 nM). One way ANOVA was performed between final polarisation values to calculate P values, and levels of significance are denoted as follows: ***P≤0.001, ns=not significant.

### Reagents

*Table S1: Plasmids used in this study, registered in the Reagents & Services database, University of Dundee (DU number)*

| NAME | PROTEIN | MUTATIONS | WT UNIPROT CODE | TAG | BACKBONE | DU NUMBER |
|---|---|---|---|---|---|---|
| **6×His-SUMO-SDE2** | SDE2 | none | Q6IQ49 | N-term 6xHis SUMO | pET15b | DU75576 |
| **HA-SDE2$_{UBL}$ (1-76)-intein-CBD** | SDE2 | Truncation (aa 1-76) | Q6IQ49 | N-term HA | pTXB1 | DU78718 |
| **Halo-SDE2-Nano** | SDE2 | none | Q6IQ49 | N-term NanoLuc, C-term HaloTag | pF Nanoluc SDE2 Halo | DU75755 |
| **USP5 WT** | USP5 | none | P45974 | N-term 6×His | pET156P | DU15641 |
| **USP5 H232A** | USP5 | H232A | P45974 | N-term 6×His | pET156P | DU78711 |
| **USP5 C335R** | USP5 | C335R | P45974 | N-term 6×His | pET156P | DU78704 |
| **USP5 M666E** | USP5 | M666E | P45974 | N-term 6×His | pET156P | DU78705 |
| **USP5 M734E** | USP5 | M734 | P45974 | N-term 6×His | pET156P | DU78706 |
| **USP5 H232A C335R** | USP5 | H232A C335R | P45974 | N-term 6×His | pET156P | DU78707 |
| **USP5 H232A M666E** | USP5 | H232A M666E | P45974 | N-term 6×His | pET156P | DU78708 |
| **USP5 H232A M743E** | USP5 | H232A M734E | P45974 | N-term 6×His | pET156P | DU78709 |
| **USP5 C3335R M666E** | USP5 | C335R M666E | P45974 | N-term 6×His | pET156P | DU78710 |
| **USP5 C3335R M743E** | USP5 | C335R M734E | P45974 | N-term 6×His | pET156P | DU78720 |
| **USP5 M666E M743E** | USP5 | M666E M734E | P45974 | N-term 6×His | pET156P | DU78712 |

*Table S2: Recombinant proteins obtained from Reagents and Services, University of Dundee*

| DU number | Construct | MW (Da) |
|---|---|---|
| **DU13025** | GST-USP2(2-353) | 72,686 |
| **DU15644** | $His_6$-USP7(1-1102) | 130,633 |
| **DU10952** | GST-USP13(318-863) | 88,475 |
| **DU14352** | $His_6$-USP14(1-494) | 58,476 |
| **DU46239** | GST-USP16(1-823) | 120,318 |
| **DU22385** | GST-USP21(196-565) | 67,325 |
| **DU20027** | Monoubiquitin | 8,559 |
| **DU20729** | Ubiquitin linear dimer | 17,640 |
| | Ubiquitin dimer (K11) | 17,640 |
| **DU20766** | Ubiquitin tetramer (linear) | 34,745 |

### Extended data

<u>Immunoprecipitation and mass spectrometry</u>

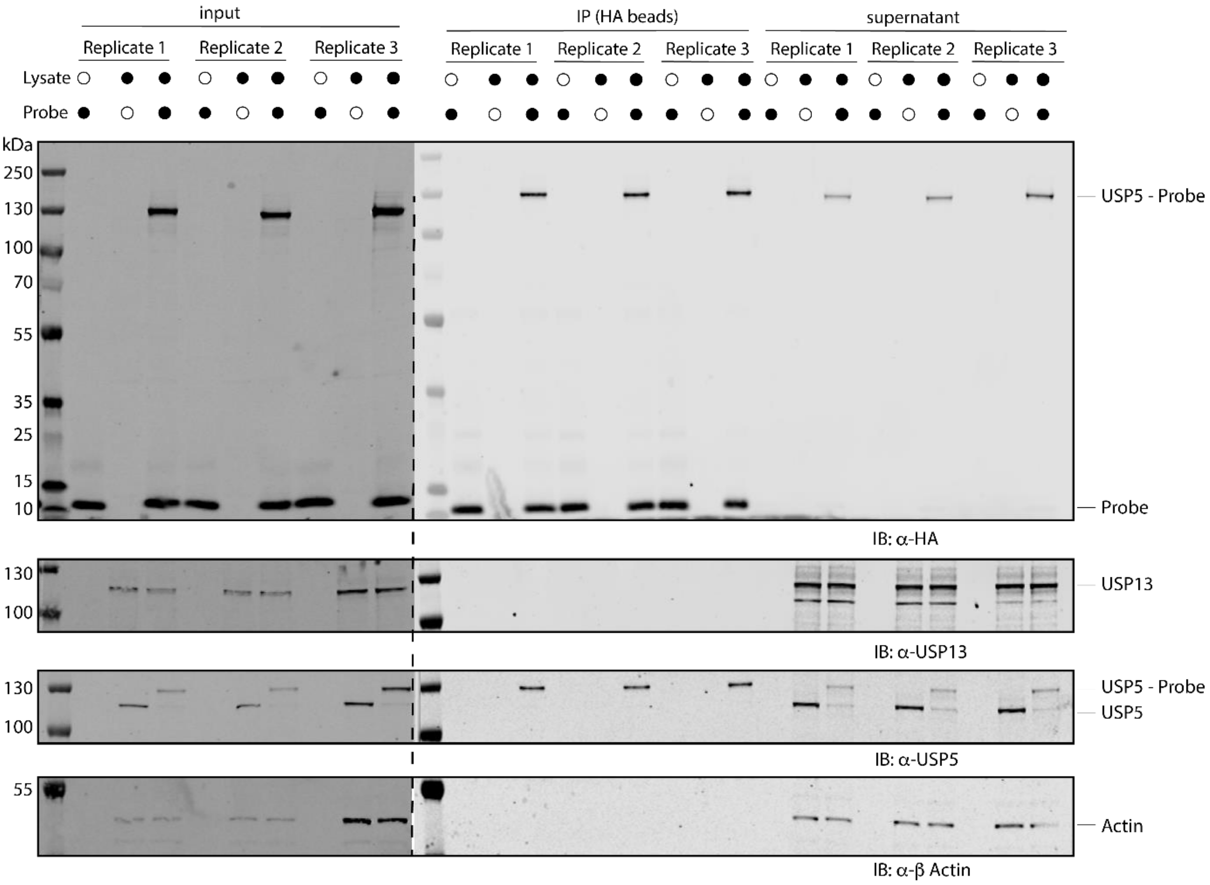


**Figure S8 | Immunoblot analysis of immunoprecipitation samples prepared for activity-based probe profiling using HA-SDE2$_{UBL}$-PA.** In triplicate, HEK293T cell lysate was incubated with HA-SDE2$_{UBL}$-PA (1 μM) for 45 min at 37 °C and immunoprecipitated with anti-HA beads. Unbound sample (supernatant) was removed and bound protein eluted (IP) by boiling.

<u>Characterisation of ABPs</u>

**HA-SDE2$_{UBL}$ΔG77-PA (HA-SDE2$_{UBL}$-PA)**

Sequence:
MYPYDVPDYAGGGGSAEAAALVWIRGPGFGCKAVRCASGRCTVRDFIHRHCQDQNVPVENF
FVKCNGALINTSDTVQHGAVYSLEPRLCG-*propargylamine*

Calculated mass (Expasy- ProtParam): 9730

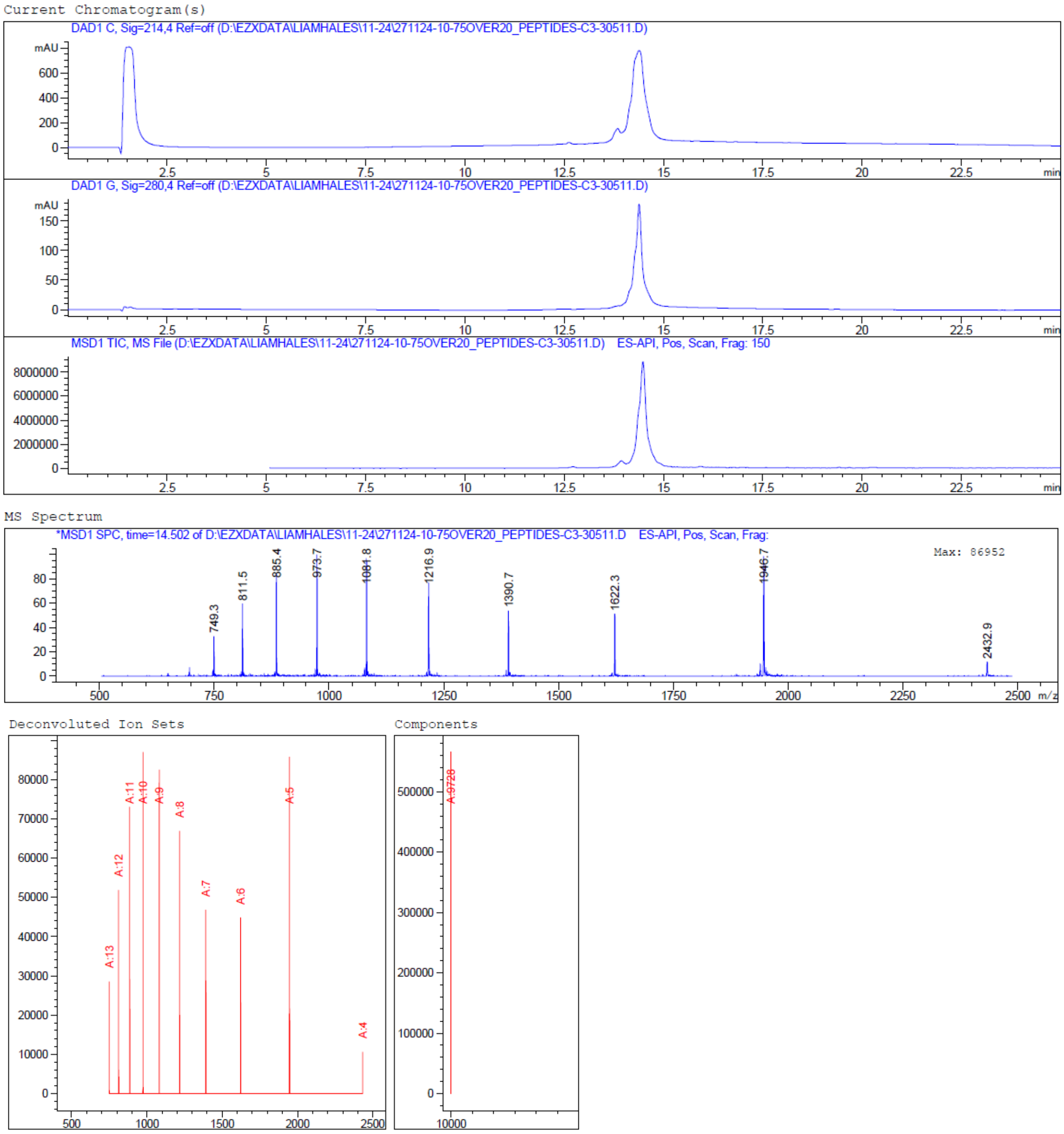

**ISG15$_{\mathrm{CTD}}$ΔG157-PA (ISG15-PA)**

Sequence:
MDEPLSILVRNNKGRSSTYEVRLTQTVAHLKQQVSGLEGVQDDLFWLTFEGKPLEDQLPLGE
YGLKPLSTVFMNLRLRG-*propargylamine*

Calculated mass (Expasy- ProtParam): 9012

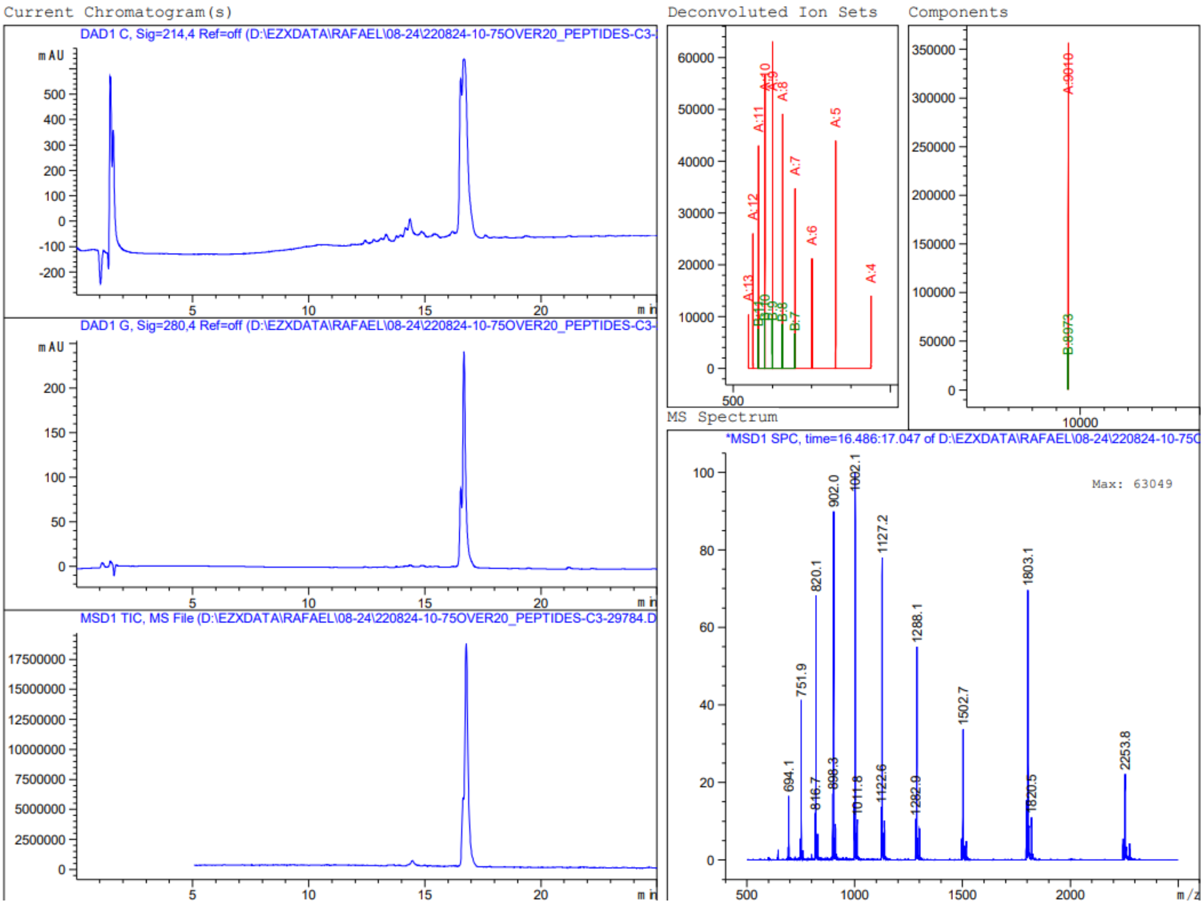

**His-UbΔG76-PA (Ub-PA)**

Sequence:
MHHHHHHHHLEVLFQGPMQIFVKTLTGKTITLEVEPSDTIENVKAKIQDKEGIPPDQQRLIFAG
KQLEDGRTLSDYNIQKESTLHLVLRLRG-*propargylamine*

Calculated mass (Expasy- ProtParam): 10,657

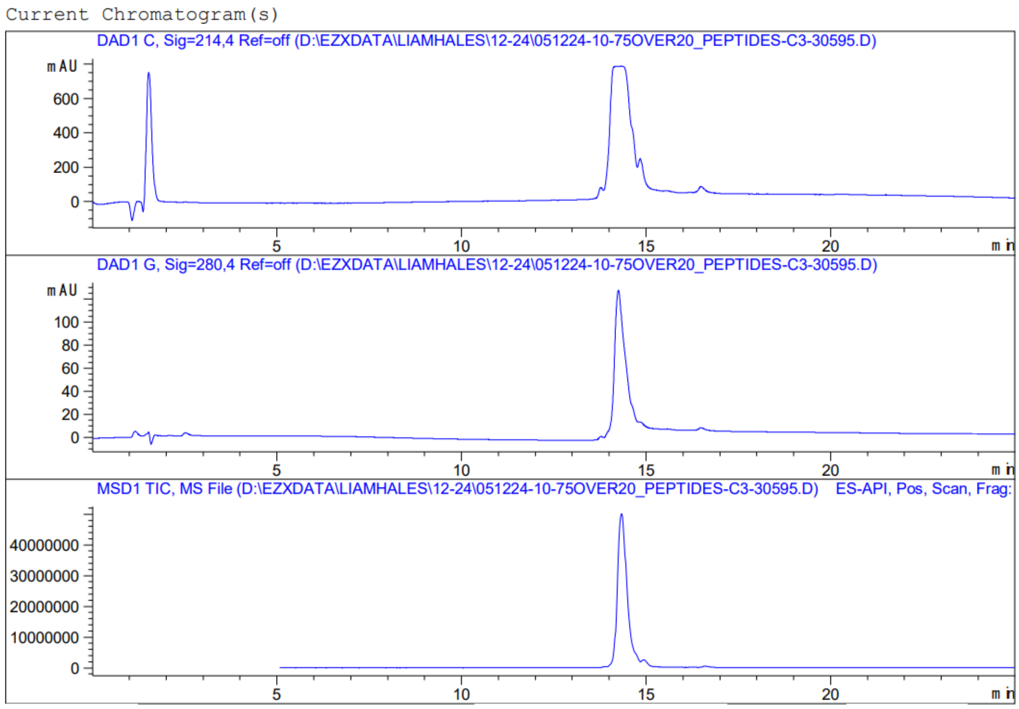


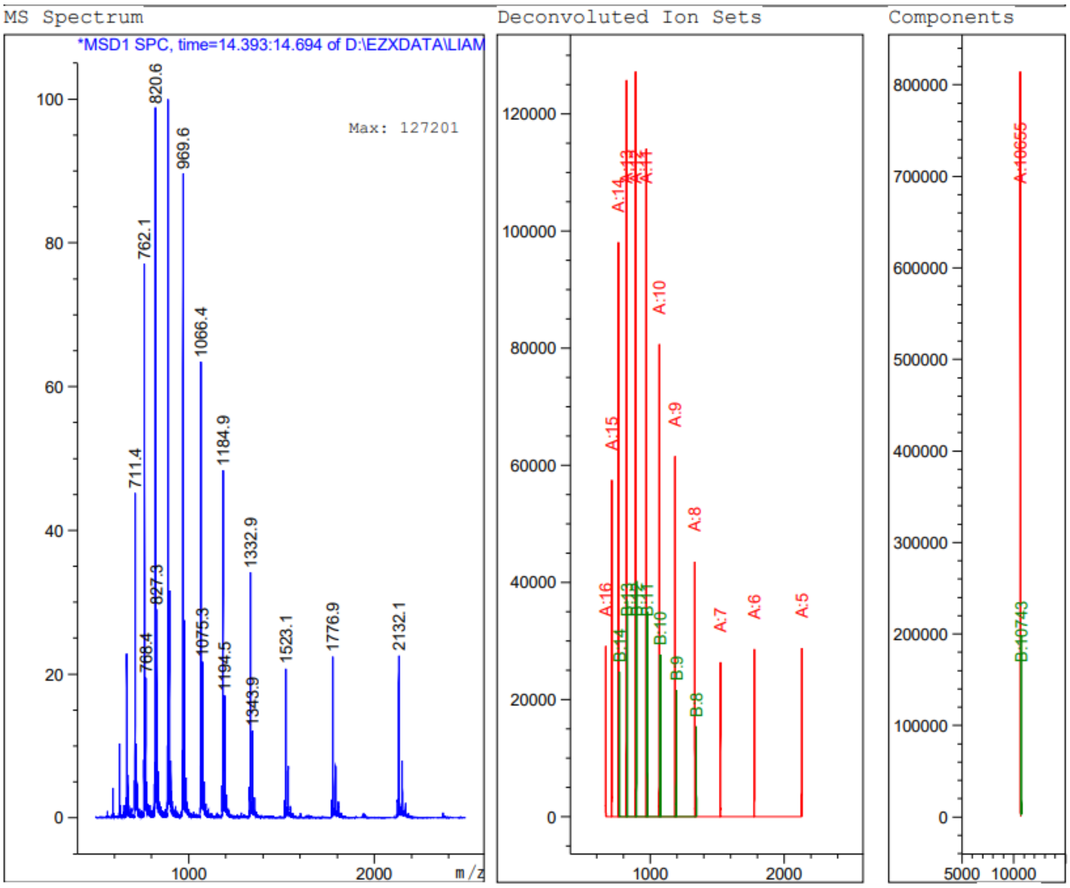

Thermal shift assay

*Table S3: Melting temperatures obtained from three technical replicates*

| | **USP5** | | | **USP5 + HA-SDE2$_{UBL}$-PA** | | | **USP5 + His-Ub-PA** | | | **USP5 + ISG15-PA** | | |
|---|---|---|---|---|---|---|---|---|---|---|---|---|
| **Melting temperature (°C)** | 53.75 | 53.75 | 54.25 | 56.25 | 56.75 | 56.25 | 58.25 | 58.75 | 58.25 | 57.25 | 57.75 | 57.25 |

ITC

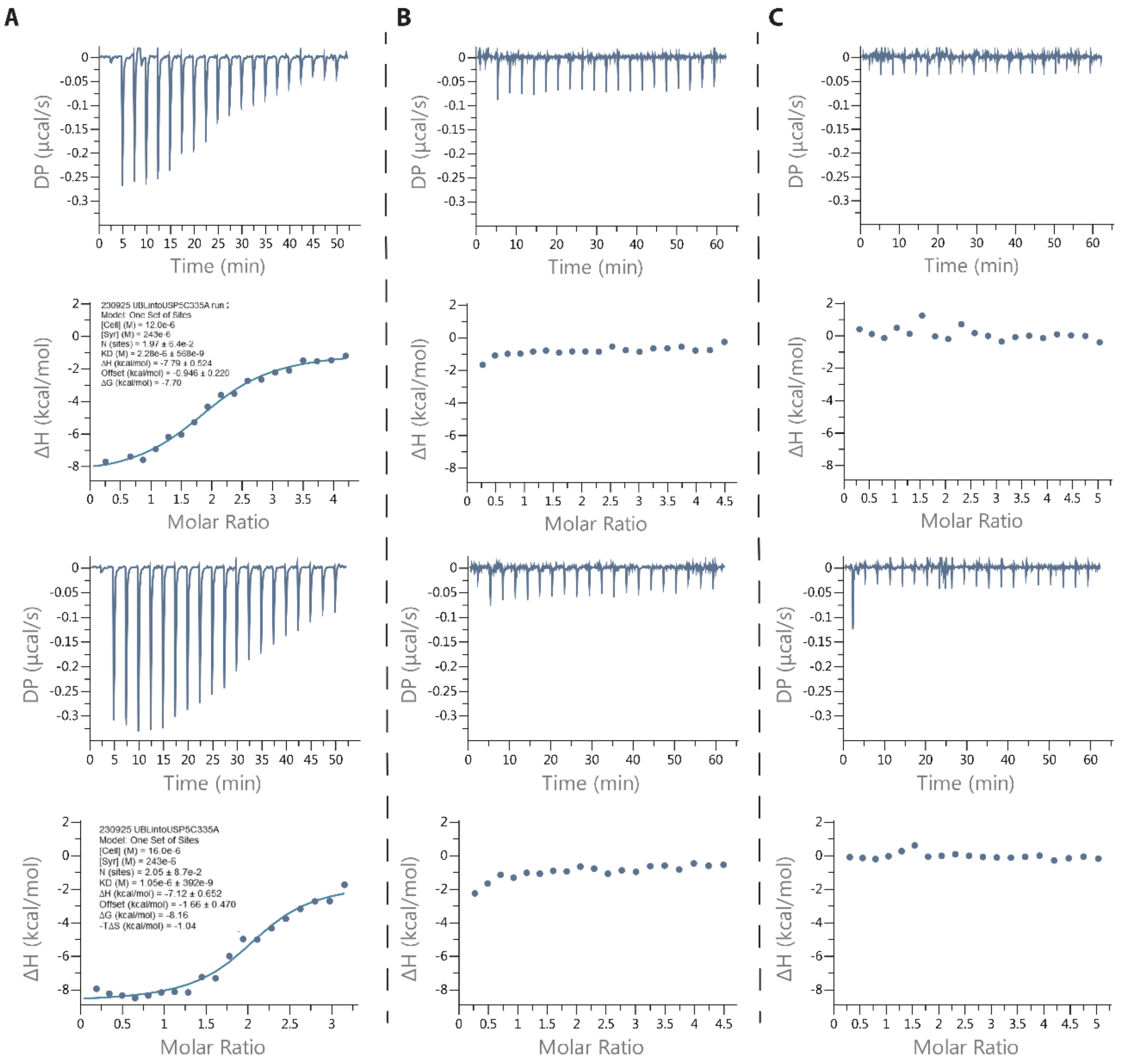


**Figure S9 | Full ITC data from the titration of SDE2 constructs into USP5 C335A**
**(A)** SDE2$_{UBL}$ (243 µM) was titrated into USP5$^{C335A}$ (12-16 µM). **(B)** SDE2$_{FL}$ (214 µM) was titrated into USP5$^{C335A}$ (10 µM). **C.** SDE2$_{CT}$ (186 µM) was titrated into USP5$^{C335A}$ (12 µM).

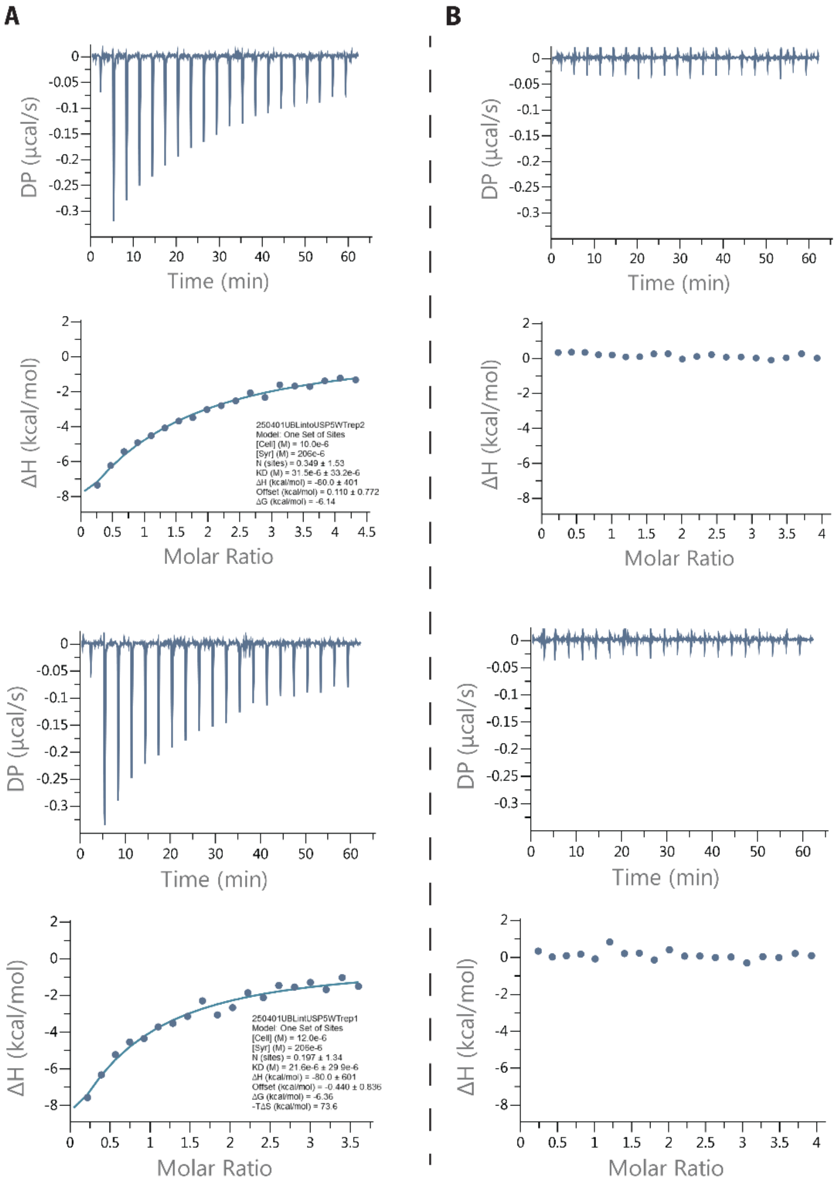


**Figure S10 | Full ITC data from the titration of SDE2$_{UBL}$ into USP5 WT and H232A**
**(A)** SDE2$_{UBL}$ (206 µM) was titrated into USP5 WT (10-12 µM). **(B)** SDE2$_{UBL}$ (206 µM) was titrated into USP5 H232A (10 µM). Experiments were performed in duplicate.

Unprocessed gels and blots

Figure 2E:

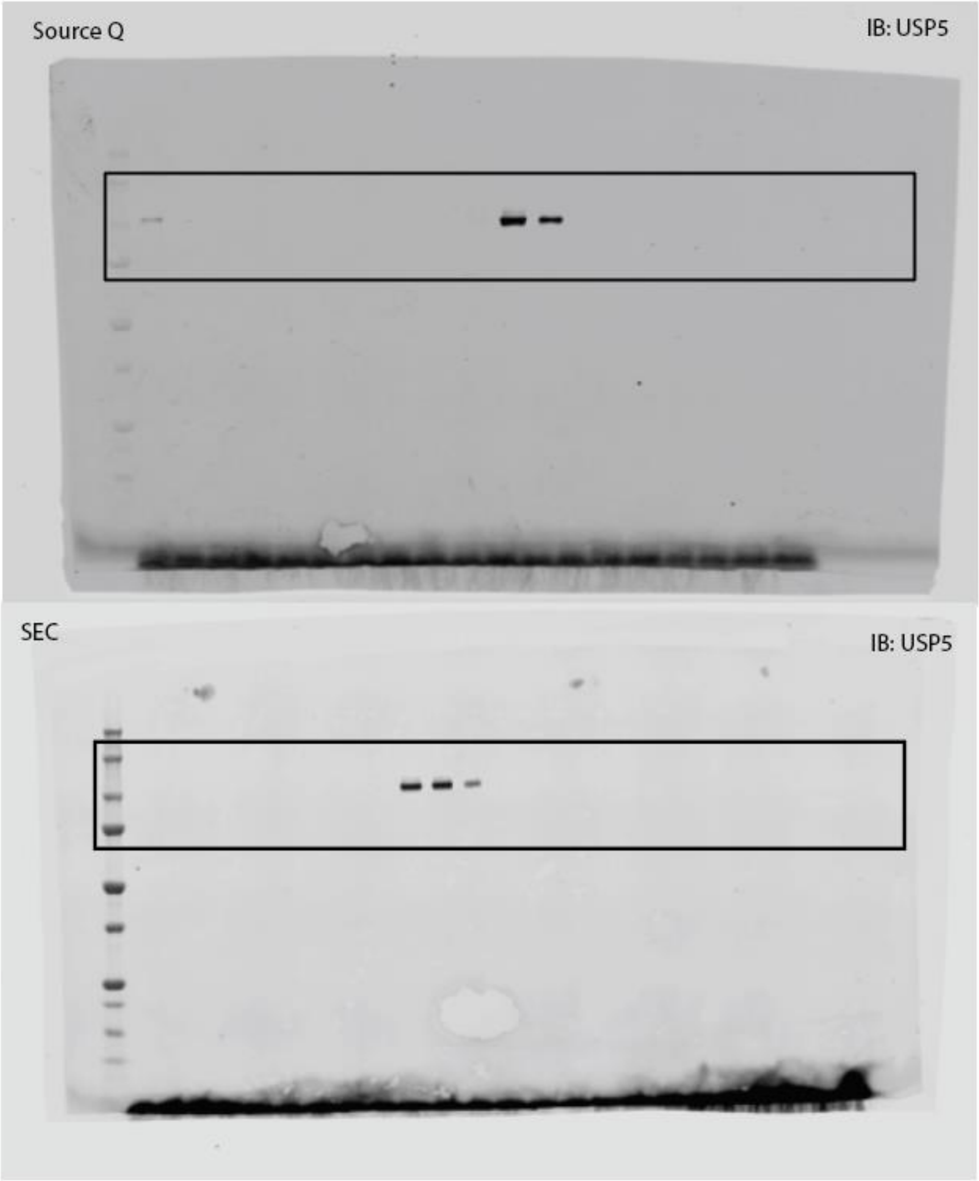

Figure 3A:

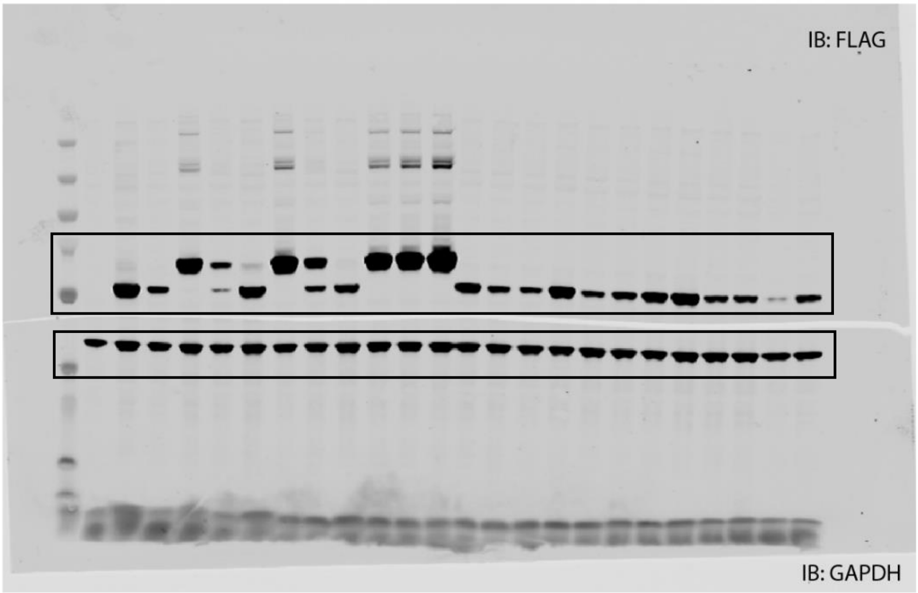


Figure 3E:

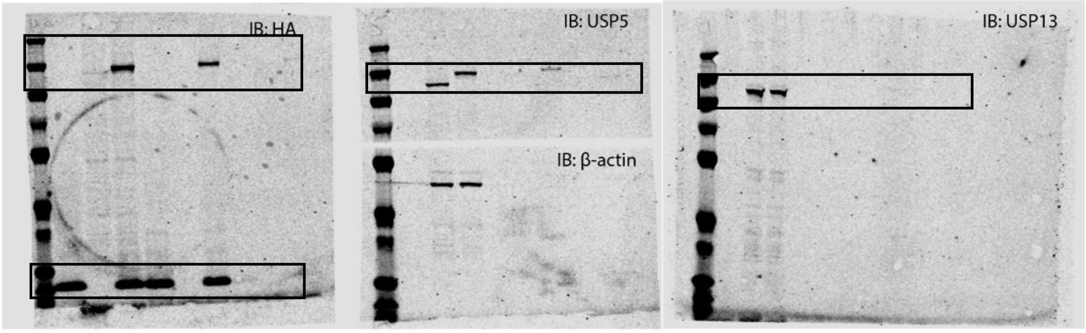

Figure 4C:

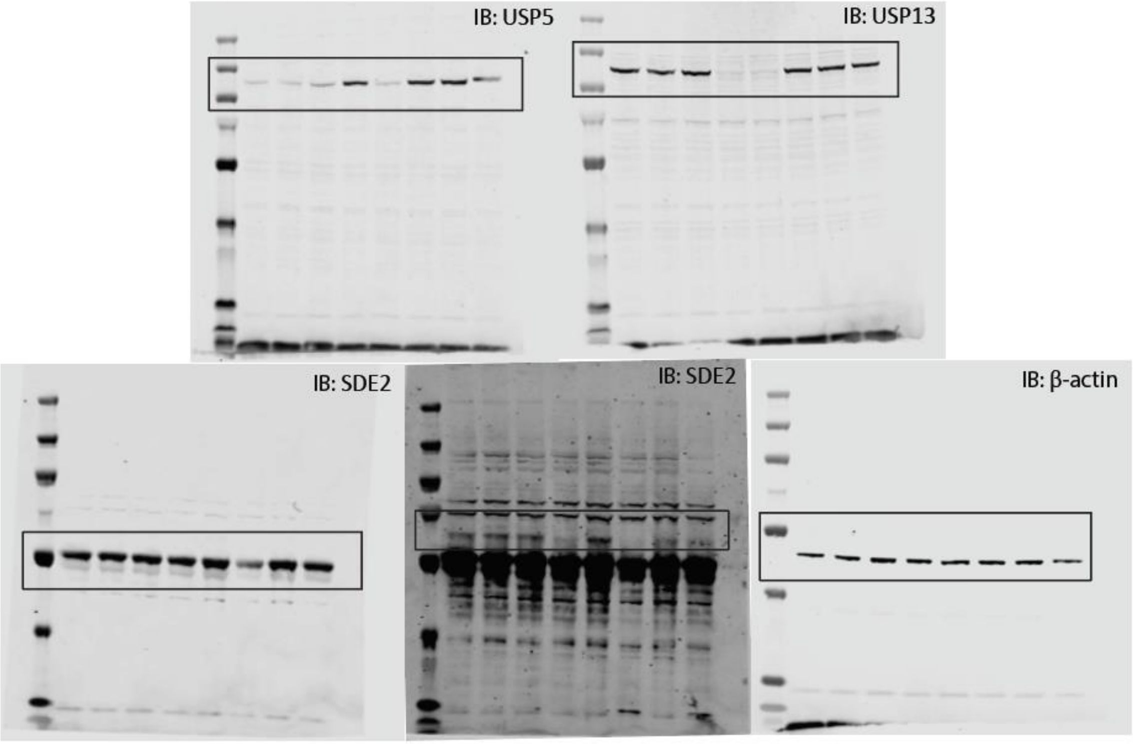


Figure S3:

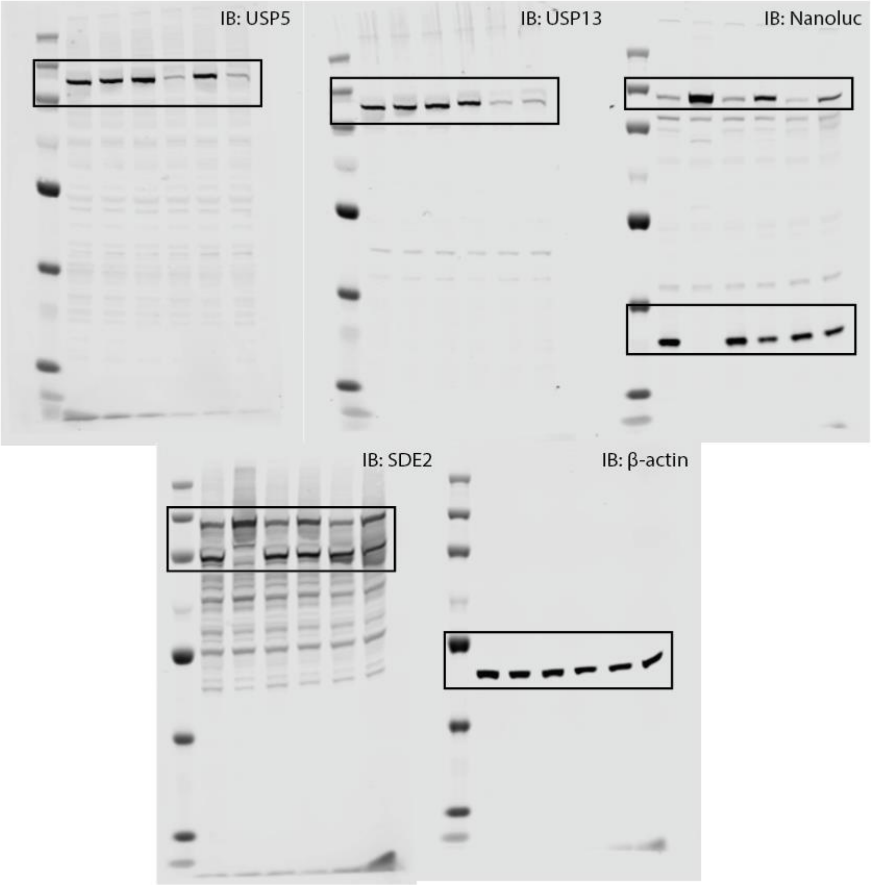

Figure 5B (Coomassie):

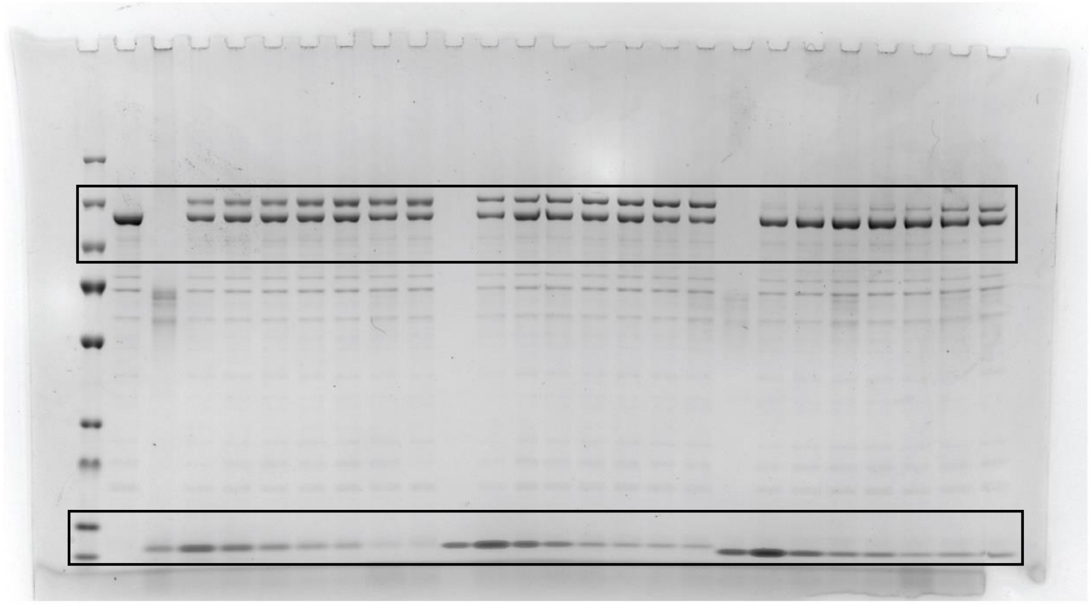

Figure 5D-F (Coomassie)

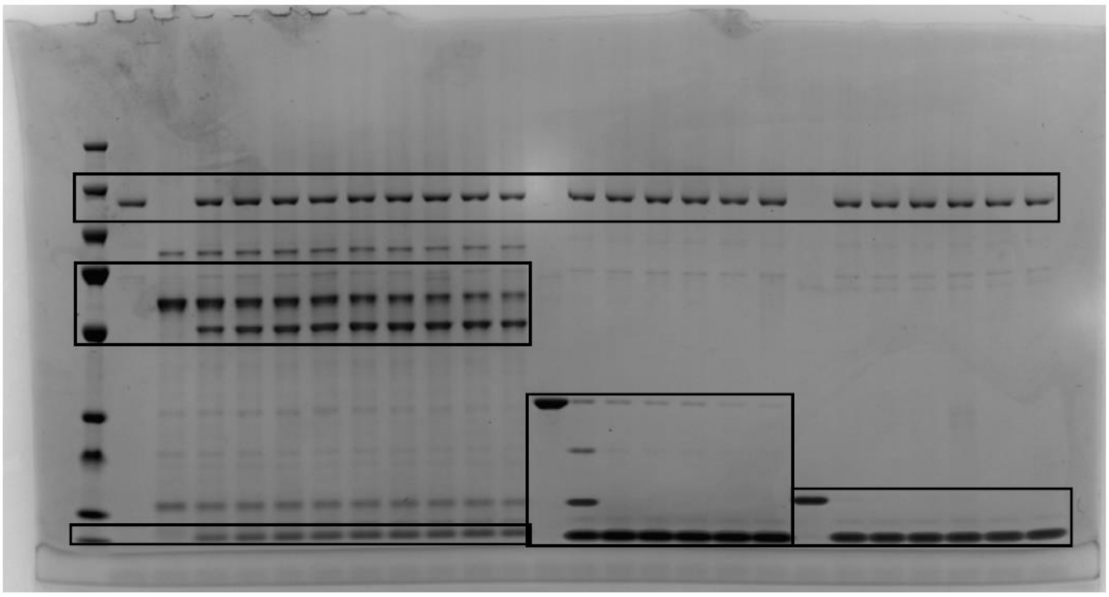
